# Ecological correlates of gene family size in a pine-feeding sawfly genome and across Hymenoptera

**DOI:** 10.1101/2021.03.14.435331

**Authors:** Kim L. Vertacnik, Danielle K. Herrig, R. Keating Godfrey, Tom Hill, Scott M. Geib, Robert L. Unckless, David R. Nelson, Catherine R. Linnen

## Abstract

A central goal in evolutionary biology is to determine the predictability of adaptive genetic changes. Despite many documented cases of convergent evolution at individual loci, little is known about the repeatability of gene family expansions and contractions. To address this void, we examined gene family evolution in the redheaded pine sawfly *Neodiprion lecontei*, a non-eusocial hymenopteran and exemplar of a pine-specialized lineage evolved from angiosperm-feeding ancestors. After assembling and annotating a draft genome, we manually annotated multiple gene families with chemosensory, detoxification, or immunity functions and characterized their genomic distributions and evolutionary history. Our results suggest that expansions of bitter gustatory receptor (GR), clan 3 cytochrome P450 (CYP3), and antimicrobial peptide (AMP) subfamilies may have contributed to pine adaptation. By contrast, there was no evidence of recent gene family contraction via pseudogenization. Next, we compared the number of genes in these same families across insect taxa that vary in diet, dietary specialization, and social behavior. In Hymenoptera, herbivory was associated with small GR and olfactory receptor (OR) families, eusociality was associated with large OR and small AMP families, and—unlike investigations in more closely related taxa—ecological specialization was not related to gene family size. Overall, our results suggest that gene families that mediate ecological interactions may expand and contract predictably in response to particular selection pressures, however, the ecological drivers and temporal pace of gene gain and loss likely varies considerably across gene families.

## Introduction

Changes in gene family size are a potentially important source of evolutionary innovation. When gene families grow via duplication, for example, reduced functional constraints may facilitate the development of phenotypic novelty (Ohno 1970; Demuth and Hahn 2009). Reductions in gene family size can also enable novel traits. For example, the colonization of highly specialized niches like oligotrophic caves (Protas et al. 2006; Gross et al. 2009; Yang et al. 2016) and toxic host plants (Matsuo et al. 2007; McBride 2007; Good et al. 2014) is linked to rampant pseudogenization. Together, these observations suggest that gene families predictably expand or contract in response to specific selection pressures. Yet compared to the rich and growing literature on genetic convergence at individual loci (Martin and Orgogozo 2013), the repeatability and predictability of gene family evolution remains understudied.

The evolution of many gene families, defined here as groups of genes that share sequence and functional similarity from common ancestry (Dayhoff 1976; Demuth and Hahn 2009), is consistent with a birth-death model where genes arise via duplication (gene gain) and are lost via deletion or pseudogenization (gene loss) (Hughes and Nei 1992; Nei and Rooney 2005). When frequency rates of duplication and deletion evolve primarily through genetic drift, over time gene family sizes contract and expand via a process dubbed genomic drift (Nei 2007; Nozawa et al. 2007). Overall, the stochastic birth-death process of genomic drift (which differs from Nei’s conceptual birth-death model of gene family evolution (Hahn et al. 2005)) sufficiently explains most gene family size distributions within genomes (Karev et al. 2002) and between species (Hahn et al. 2007).

But during an ecological shift, natural selection can influence birth-death dynamics by promoting the expansion or contraction of specific gene families. Thus, taxa adapted to a novel niche may have genomic evidence of selective maintenance for gene duplications or deletions. For example, if selection favors gene gain, novel gene duplicates will tend to persist in the population and form subfamilies of recently diverged paralogs. If the mutational mechanism that generates new duplicate genes is unequal crossing over during meiosis, these recently diverged paralogs will be arranged in tandem arrays across the genome (Zhang 2003). Moreover, if duplicates experience positive selection for novel functions, they can have elevated amino acid substitution rates. Conversely, some genetic functions may become obsolete or even deleterious in the novel habitat. In this case, positive or relaxed purifying selection will cause some gene families to accumulate loss-of-function mutations at an accelerated rate.

After an ecological shift, impacted gene families will eventually reach a new equilibrium state where gene number evolves primarily through negative selection and genomic drift. Likewise, tandem array lengths will reflect local recombination rates (Akhunov et al. 2003; Zhang and Gaut 2003; Rizzon et al. 2006; Thomas 2006) and pseudogenes will fade into the genomic background (Petrov et al. 1996; Petrov and Hartl 1997, 1998). Thus, within-genome signatures of adaptive changes in gene family size are likely ephemeral and best detected in lineages that *recently* shifted to a novel niche. Plus, if selection consistently favors the expansion or contraction of specific gene families in specific environments, among-taxon correlations between gene family size and ecology should be maintained. Currently, the extent to which different taxa converge at the level of gene family size changes is largely unknown.

Arguably, the genes most likely to expand and contract convergently in response to similar selection pressures are those that mediate organismal interactions with their biotic and abiotic environments. These “environmentally responsive genes” include chemosensory (e.g., olfactory and gustatory receptors), detoxifying (e.g., cytochrome P450), and immunity (e.g., immunoglobulin and MHC) genes. To cope with constantly changing pressures, environmentally responsive genes tend to be characterized by elevated sequence diversity, duplication rates, substitution rates, and genomic clustering, as well as tissue- or temporal-specific expression (Berenbaum 2002) and limited pleiotropy (Arguello et al. 2016). Importantly, causal links between changes in environmentally responsive genes and adaptation to novel niches have been established for multiple taxa (Després et al. 2007; Matsuo et al. 2007; Dobler et al. 2012; Zhen et al. 2012; Sezutsu et al. 2013).

With exceptionally diverse ecologies and an ever-increasing availability of annotated genomes (Consortium 2013; Poelchau et al. 2015), insects are a powerful system for investigating the predictability of size changes in environmentally responsive gene families. To date, at least two ecological transitions are hypothesized to have a predictable impact on gene family size in insect lineages. In plant-feeding insects, the evolution of increased dietary specialization (i.e., smaller diet breadth) is associated with reduced chemosensory and detoxifying gene family sizes (McBride 2007; McBride and Arguello 2007; Good et al. 2014; Goldman-Huertas et al. 2015; Calla et al. 2017; Comeault et al. 2017) but see (Gardiner et al. 2008). In hymenopteran insects, eusociality is associated with expansions of the olfactory-receptor family and contractions of the gustatory-receptor family (Robertson and Wanner 2006; Zhou et al. 2015; McKenzie et al. 2016; Brand and Ramírez 2017) but see (Fischman et al. 2011; Johnson et al. 2018). Most of these studies, however, consider a single ecological characteristic or gene family (but see Robertson and Wanner 2006) which is problematic since changes in social behavior may often be accompanied by changes in other ecological characteristics and vice versa (Faulkes et al. 1997; Duffy and Macdonald 2010; Ross et al. 2013). A better understanding of ecology and gene family size relationships requires simultaneous consideration of multiple ecological characteristics and diverse gene families.

Here, we characterize multiple environmentally responsive gene families in the genome of the redheaded pine sawfly, *Neodiprion lecontei* (Order: Hymenoptera; Family: Diprionidae). This species provides an opportunity to examine both within-genome signatures of adaptive gene family contractions and expansions, and among-lineage correlations between ecology and gene family size. First, for within-genome signatures, *N. lecontei* is an exemplar of an herbivorous hymenopteran lineage (Diprionidae) that underwent a drastic host shift: sometime within the last 60 million years, this lineage transitioned from angiosperms to coniferous host plants in the family Pinaceae (Boevé et al. 2013; Peters et al. 2017). To defend against herbivores and pathogens, Pinaceae produce viscous oleoresin secretions that are sticky and have unique antimicrobial properties (Trapp and Croteau 2001; Gershenzon and Dudareva 2007). To manage these toxic and extraordinarily sticky resins, *N. lecontei* and related diprionids evolved specialized feeding and egg-laying traits (Figure 1). Beyond these traits, we hypothesize that pine specialization likely resulted in pronounced changes to the selection pressures acting on multiple gene families, especially those involved in chemosensation, detoxification, and immune function. Second, with respect to among-lineage correlations between ecology and gene-family size, *N. lecontei* is an herbivorous, non-eusocial insect from the Eusymphyta, a massively understudied hymenopteran clade (Peters et al. 2017). Although many assembled and annotated hymenopteran genomes are currently available, almost all have come from apocritans (bees, wasps, and ants, but see (Robertson et al. 2018)). Thus, *N. lecontei* increases the ecological, behavioral, and taxonomic diversity of hymenopteran genomes for evaluating ecological correlates of gene family size among taxa.

**Figure 1.**
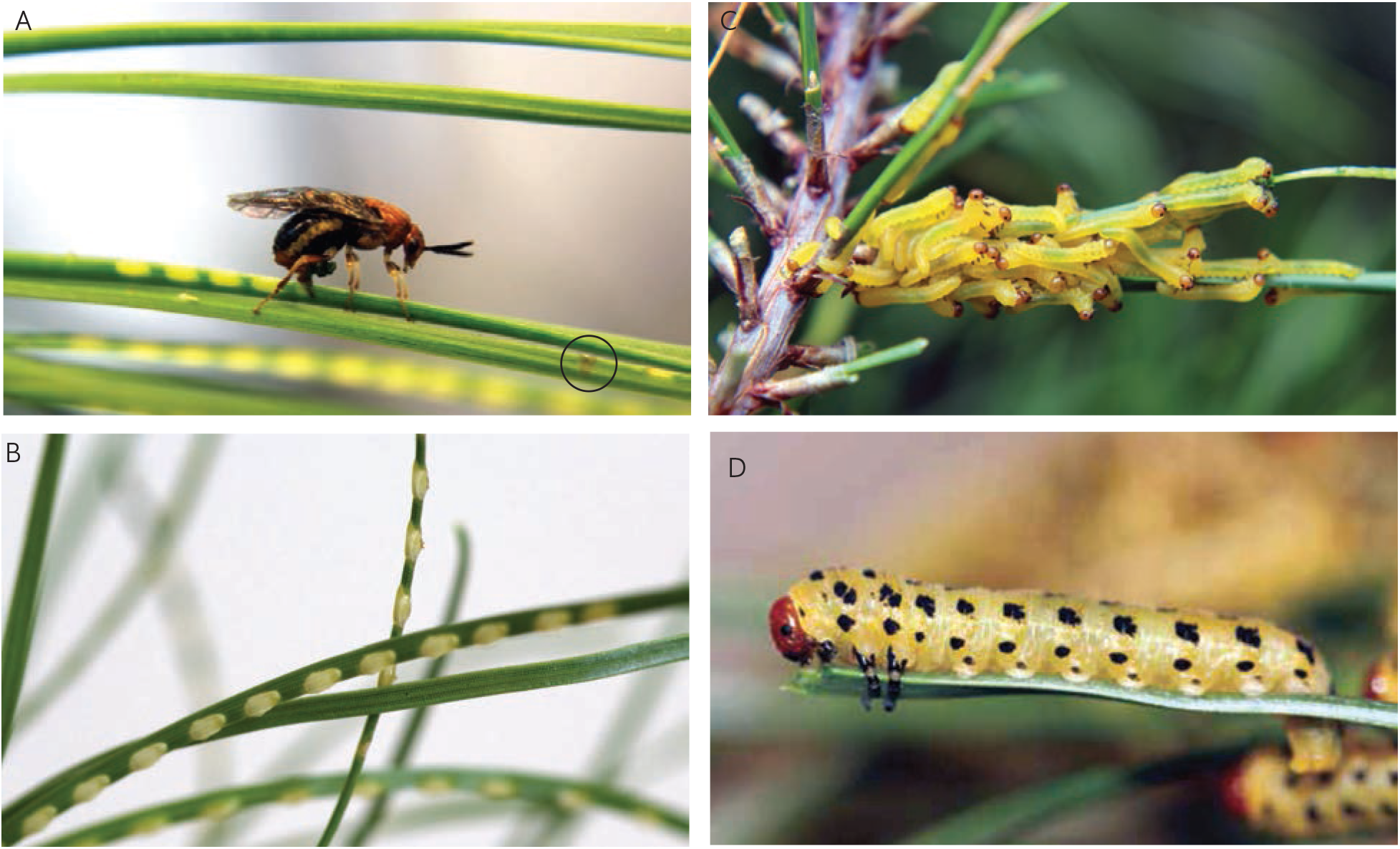
Like other diprionids, *N. lecontei* has multiple morphological and behavioral adaptations to *Pinus* foliage. **A.** An egg-laying *N. lecontei* female demonstrating several adaptations for dealing with thick, resinous pine needles, including: a robust saw-like ovipositor (visible within the needle), a tendency to lay many closely spaced eggs per needles, and a tendency to cut resin-draining slits on egg-bearing needles (circled). **B.** Prior to hatching, *N. lecontei* eggs absorb water from the host, causing the eggs to swell and the pockets to open. Throughout development, embryos are in close contact with living host tissue. **C.** Early-instar larvae have skeletonizing feeding behavior in which only the outer needle tissue is consumed, leaving the resinous interior intact. This strategy prevents small larvae from being overwhelmed by sticky resin. **D.** Mid- and late-instar larvae consume the entire pine needle. Larvae sequester pine resin in specialized pouches for use in self-defense (All photos by R.K. Bagley).

To evaluate the predictability of gene family evolution, we assembled a draft genome for *N. lecontei* and manually annotated genes for five environmentally responsive gene families: olfactory receptor (OR), gustatory receptor (GR), odorant binding protein (OBP), cytochrome P450 (CYP), and antimicrobial peptide (AMP). For gene families that underwent a size change related to pine adaptation, we expected one or more of the following patterns: (1) clusters of recently diverged paralogs in gene-family trees, (2) a high proportion of genes in tandem arrays, (3) signatures of positive selection among paralogs, and (4) elevated rates of pseudogenization. Then, for the same five gene families, we asked whether gene-family size correlated with ecology among distantly related insect taxa. To do so, we compiled published gene annotations and ecological variables (diet type, degree of ecological specialization, presence/absence of eusociality) for hymenopteran taxa. Together, these analyses identify possible candidate gene families underlying pine specialization and reveal that relationships between gene family size and ecology differ among environmentally responsive gene families.

## Results

### Genome assembly and annotation

#### Sequencing and assembly

We sequenced one mate-pair and two small-insert Illumina libraries made from haploid male siblings (see Methods). After read processing, we retained 268 billion PE100 reads with a combined read depth of 112x (Table S1). ALLPATHS-LG (v47417) (Gnerre et al. 2011) produced a 239-Mbp assembly consisting of 4523 scaffolds, with a scaffold N50 of 243 kbp (Table S2). Prior studies identified seven chromosomes in *N. lecontei* (Smith 1941; Maxwell 1958; Sohi and Ennis 1981; Linnen et al. 2018). With an estimated genome size (1C) of 331 ±9.6 Mbp, our assembly captured 72% of the genome. Overall, these metrics are comparable to other hymenopteran assemblies (Table S2).

To measure assembly completeness and artificial sequence duplication, we used CEGMA (Parra et al. 2007) and BUSCO (Simão et al. 2015). Both search the assembly for a set of single-copy, conserved genes, however, the CEGMA software has been deprecated (http://korflab.ucdavis.edu/Datasets/cegma). Of the 248 CEGMA core eukaryotic genes, 90% aligned as complete, single copies and 8% aligned complete but duplicated. For BUSCO, we used the OrthoDB arthropod dataset, and out of 2675 groups 77% were complete, single copies and 3% were complete but duplicated. These metrics indicate the presence of artificial duplicate sequences, but otherwise the assembly was reasonably complete and suitable for annotation.

About 15.8% of the assembly consisted of repetitive elements, including 122 unknown transposable elements that were mostly unique to *N. lecontei* (Table S3), and 212 other families of transposable elements and simple repeats. This 15.8% corresponds to 11.4% of the actual 331-Mb genome, of which we predict 27.6% is repetitive, suggesting that ~16.1% of the missing ~28% of the genome is repetitive content (Table S3). For de novo gene prediction, we included the *N. lecontei* transcriptome and protein-coding genes from *Atta cephalotes* (OGSv1.2), *Acromyrmex echinatior* (OGSv3.8), *Apis melifera* (OGSv3.2), *Athalia rosae* (OGSv1.0), and *Nasonia vitripennis* (OGSv1.0) to guide annotation. The official gene set (OGSv1) had 12,980 gene models while the transcriptome had an average of 26,000 transcripts per tissue (Table S4).

#### Olfactory receptor

The OR gene family had 56 genes total, including the co-receptor *Orco*; one gene contained stop codons, three were partial annotations, and 52 genes were intact (Table 1). In *D. melanogaster* most olfactory sensory neurons (OSNs) express a single OR (along with the coreceptor, *Orco*), and OSNs expressing a particular OR converge on a single glomerulus in the antennal lobe (Gao et al. 2000; Vosshall et al. 2000; Couto et al. 2005) but see (Fishilevich and Vosshall 2005). This anatomy results in a general one-to-one correspondence between the number of ORs and the number of glomeruli, a correspondence also observed in the hymenopteran European honey bee (*Apis mellifera*, (Robertson and Wanner 2006)). Based on these studies and examination of the antennal lobes of adult male and adult female *N. lecontei*, we expected to find a minimum of 49 functional ORs (Table S5, Figure 2). The close correspondence between our gene annotations and glomeruli counts suggests that we have located all or most *N. lecontei* OR genes.

**Table 1.** Summary of within-genome signatures of adaptive expansions and contractions for five environmentally responsive gene families.

|  | Gene family size |  |  |  |  | Genomic Clustering |  | Molecular evolution |  |  |
| --- | --- | --- | --- | --- | --- | --- | --- | --- | --- | --- |
| Gene family* | Intact genes | Partial | Pseudo | Total genes | Prop. pseudo | Prop. in clusters<sup>†</sup> | Largest cluster | *Neodiprion*-specific clades<sup>‡</sup> | Significant branch tests<sup>§</sup> | Significant site tests** |
| OR | 52 | 3 | 1 | 56 | 0.02 | 0.59 | 8 | 3 | 1 | 0 |
| GR | 41 | 2 | 2 | 44<sup>††</sup> | 0.05 | 0.76 | 10 | 3 | 1 | 1 |
| OBP | 13 | 0 | 0 | 13 | 0 | 0.38 | 3 | 0 | n/a | n/a |
| CYP (all) | 93 | 2 | 12 | 107 | 0.11 | 0.66 | 16 | 5 | 2 | 0 |
| CYP2 clan | 9 | 0 | 0 | 9 | 0 | 0.33 | 2 | 0 | 0 | 0 |
| CYP3 clan | 47 | 0 | 8 | 55 | 0.15 | 0.81 | 16 | 4 | 2 | 0 |
| CYP4 clan | 27 | 2 | 4 | 33 | 0.12 | 0.55 | 3 | 1 | 0 | 0 |
| mito CYP clan | 10 | 0 | 0 | 10 | 0 | 0.50 | 3 | 0 | 0 | 0 |
| AMP | 21 | 0 | 0 | 21 | 0 | 0.95 | 15 | ?<sup>‡‡</sup> | 0 | 0 |

\* Abbreviations: OR = olfactory receptor genes; GR = gustatory receptor genes; OBP = odorant binding protein genes; CYP = cytochrome P450 genes (“clans” refer to four major clades of CYPs present in insects); AMP = antimicrobial peptide genes.
<sup>†</sup> Calculated as: (number of genes in clusters of 2 or more)/(genes for which clustering could be evaluated).
<sup>‡</sup> Defined as monophyletic clusters of 5 or more *Neodiprion* paralogs with a bootstrap support $\geq 70\%$ in an amino acid phylogeny constructed with gene annotations from *Neodiprion*, select Hymenoptera, and *Drosophila melanogaster*.
<sup>§</sup> To be counted, clades had to reject both 1-ratio and fixed-ratio models in dN/dS branch tests (see Table 2).
\*\* To be counted, clades had to reject both M7 and M8a models in dN/dS site tests (see Table 2).
<sup>††</sup> One gene was both a partial annotation and a pseudogene.
<sup>‡‡</sup> Low bootstrap support precluded the identification of *Neodiprion*-specific clades.

**Figure 2.**
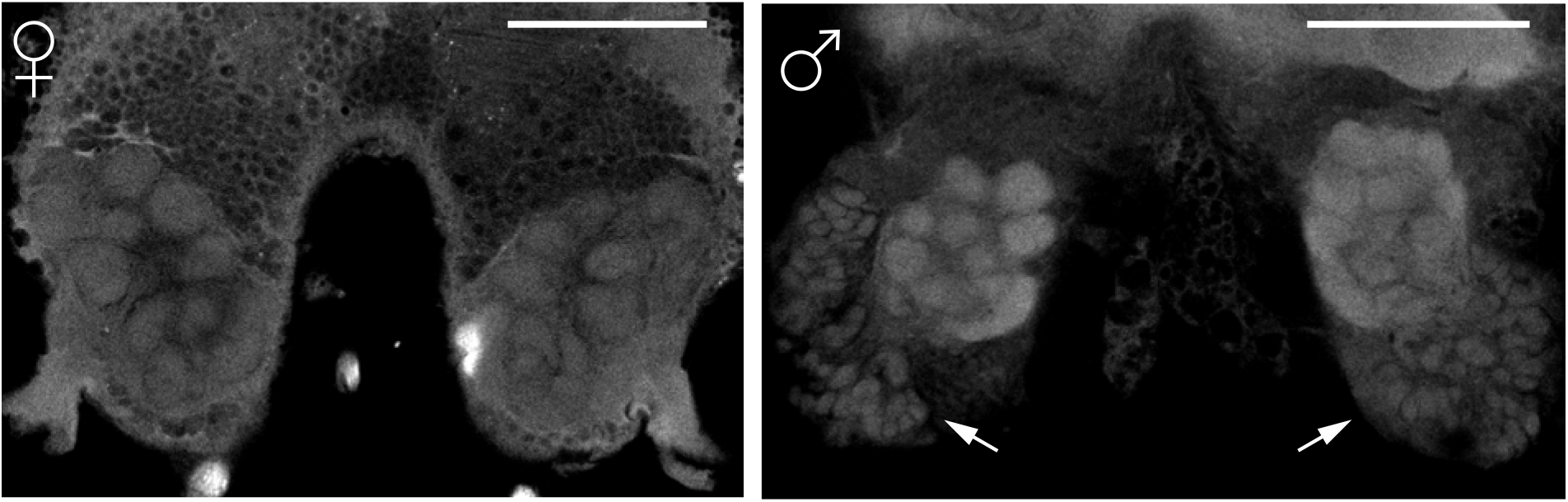
Optical sections through the antennal lobes of adult female (left) and male (right) *N. lecontei*. White arrows indicate regions of male-specific synaptic clusters. Scale bars = 500 μm.

59% of ORs were in genomic clusters of two or more genes (Figure 3), a low proportion compared to many other hymenopteran OR families (Robertson and Wanner 2006; Zhou et al. 2015; McKenzie et al. 2016; Brand and Ramírez 2017). A phylogenetic analysis of OR protein sequences from *Neodiprion,* six other hymenopterans, and *D. melanogaster* identified three *Neodiprion*-specific clades with at least five genes (Figure S1a). These same three clades were also recovered in a phylogenetic analysis of *Neodiprion* OR cDNA sequences (Figure S1b). For each *Neodiprion*-specific OR clade (and *Neodiprion*-specific clades in other gene families, see below), we used the *Neodiprion* cDNA tree, the codeml program in the PAML package (Yang 2007), and likelihood-ratio tests to ask: (1) whether the ratio of non-synonymous to synonymous substitution rates (dN/dS or ω) for the focal OR clade differed from the rest of the *Neodiprion* OR gene family and, if so, whether they exhibited evidence of positive selection (ω>1) (branch tests); and (2) whether ω differed among sites across members of *Neodiprion*-specific clades and, if so, which sites exhibited evidence of positive selection (site tests). For only one OR clade (OR clade 1) did we detect evidence of branch-specific positive selection (i.e., rejection of both 1-ratio and fixed-ω models), but this clade lacked evidence of site-specific positive selection (Table 2).

**Figure 3.**
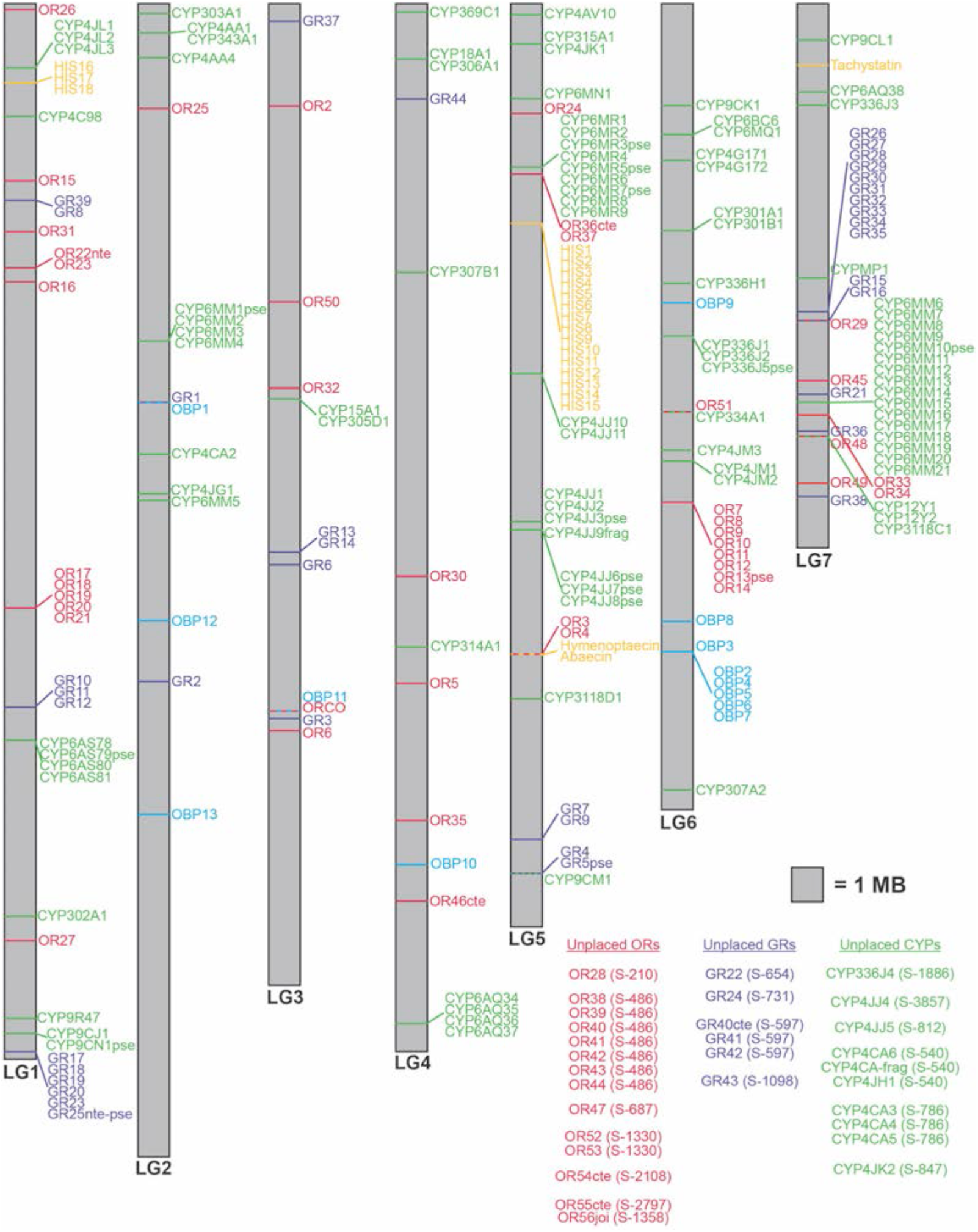
Position of genes belonging to five environmentally responsive gene families along seven *N. lecontei* linkage groups. Linkage groups (LG) are drawn to scale and ordered as in the linkage-group anchored assembly described in Linnen et al. 2018 (GenBank accession numbers are as follows: LG1 = CM009916.1; LG2 = CM009917.1; LG3 = CM009918.1; LG4 = CM009919.1; LG5 = CM009920.1; LG6 = CM009921.1; LG7 = CM009922.1). Gene family abbreviations: OR (olfactory receptor), GR (gustatory receptor), OBP (odorant binding protein), CYP (cytochrome P450), AMP (antimicrobial protein). Each gene family is represented by a different color. Horizontal lines indicate the approximate locations of genes within LG; diagonal lines that connect to horizontal lines are used to highlight groups of genes that met our clustering criteria. Genes that were found on scaffolds that have not been placed on linkage groups are indicated on the bottom left, with abbreviated scaffold names given in parentheses (e.g., S-210 = scaffold_210 = LGIB01000210.1 in the assemblies available on NCBI).

**Table 2.**
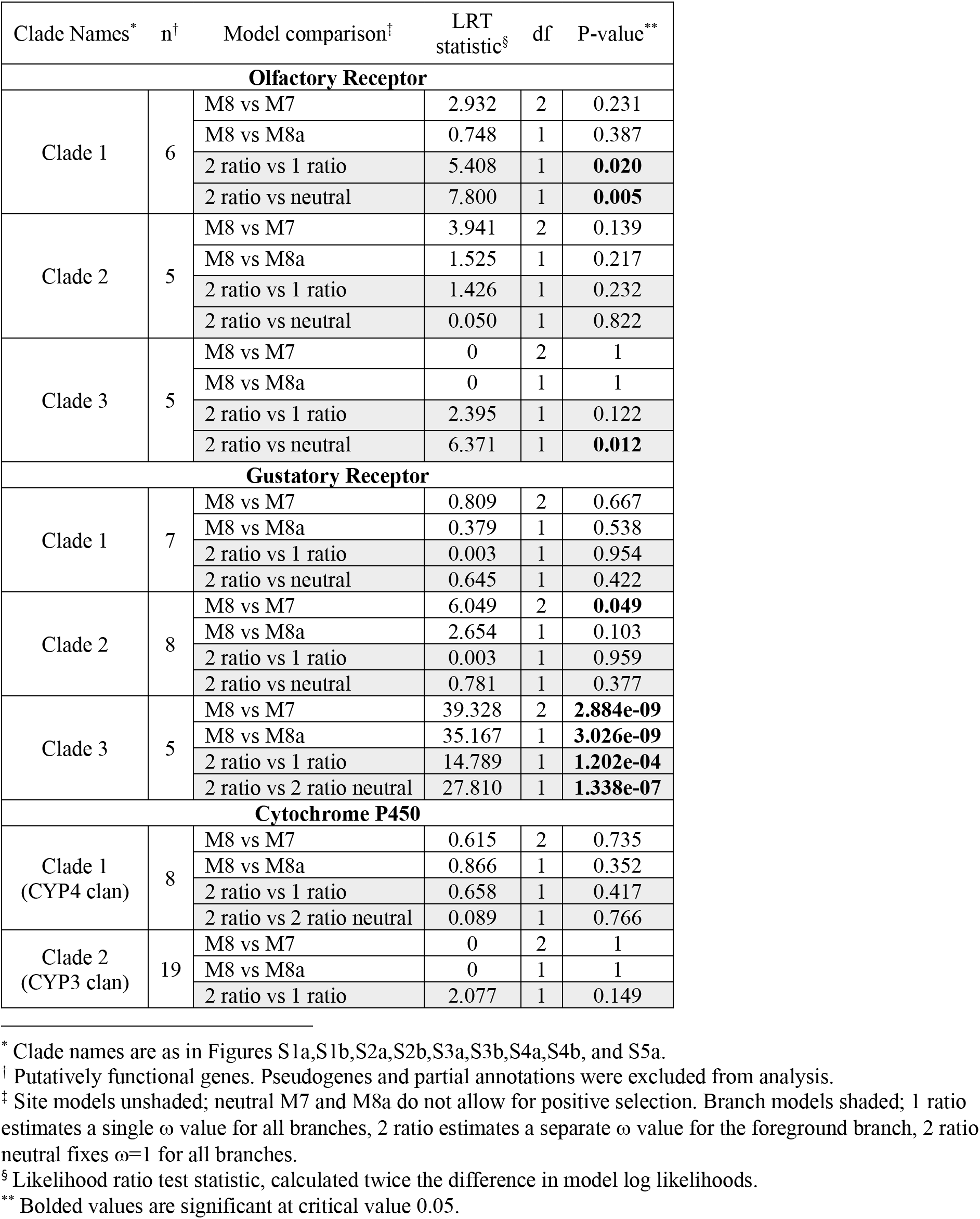
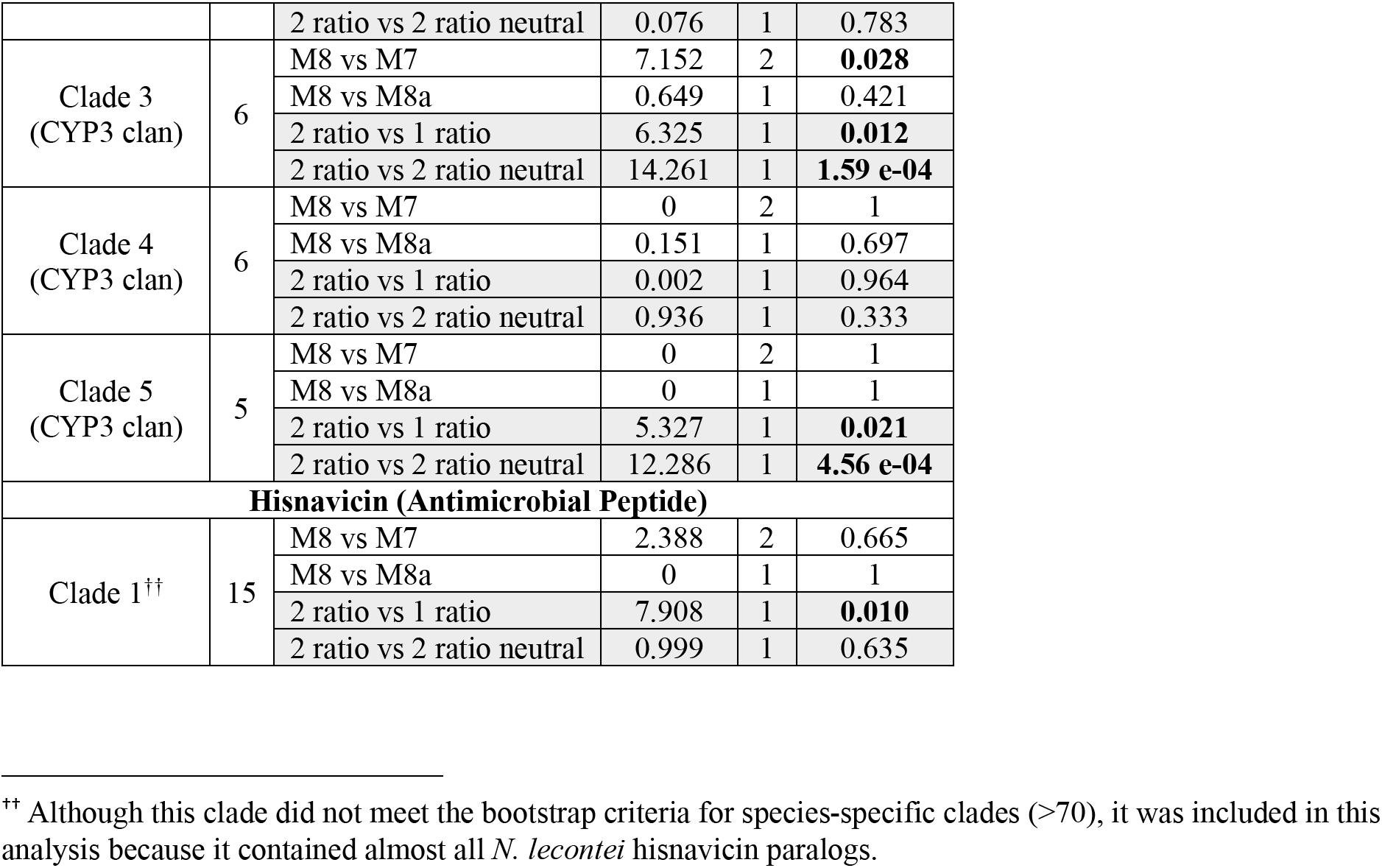
Likelihood-ratio tests (LRTs) of positive selection on *Neodiprion*-specific clades (branch models) and on amino acid sites within these clades (site models).

#### Gustatory receptor

The GR gene family had 44 genes total; two genes contained stop codons, two were partial annotations (one annotation was both partial and pseudogenized), and 41 were intact (Table 1). 76% of the GRs that could be placed on chromosomes were in genomic clusters (Figure 3) with three *Neodiprion*-specific clades of at least five genes (Figures S2a and S2b). Only one clade (GR clade 3) had evidence of branch-specific positive selection (Table 2). This clade also had evidence of positive selection at some amino acid positions among paralogs (Table 2; sites with evidence of positive selection include: 77E, 79S, 146N, 275S, 301S). Notably, GR Clade 3 is an expansion of six paralogs orthologous to *DmGR66a*, a bitter receptor specifically for caffeine (Moon et al. 2006). However, *N. lecontei* orthologs were not found for *DmGR93a* (Lee et al. 2009) and *DmGr33a* (Moon et al. 2009), coreceptors possibly required for caffeine detection. Together, these data suggest that caffeine-like GR receptors have been coopted for novel functions in *N. lecontei*.

The GR family also had orthologs for sugar receptors *DmGR5a* (trehalose) (Dahanukar et al. 2001), *DmGR43a* (fructose) (Miyamoto et al. 2012), and *DmGR64a-f* (multiple sugars) (Slone et al. 2007) as well as carbon dioxide receptors *DmGR21a* and *DmGR63a* (Jones et al. 2007) (Figure S2a). Orthologs to these carbon dioxide receptors have not been found in Apocrita but seem to be preserved in Symphyta, like *N. lecontei* (Robertson and Kent 2009; Robertson et al. 2018).

#### Odorant binding protein

The OBP gene family had 13 genes total; none were pseudogenized or partial annotations (Table 1). In this family, 38% of genes were in genomic clusters, including a cluster of five genes on chromosome 6 (Figure 3). *Neodiprion*-specific OBP clades were not found, even for the chromosome 6 cluster. We note, however, that the OBP phylogenies had low bootstrap support (Figure S3a,b), making it difficult to infer relationships among paralogs.

#### Cytochrome P450

The CYP gene family had 107 genes total; twelve genes contained stop codons, two were partial annotations, and 93 were intact (Table 1). In insects, CYPs belong to four major clades, which are referred to as clans (Feyereisen 2012). When we split the CYP gene family by clan, the CYP2 clan had nine intact genes; the CYP3 clan had 47 intact genes and eight pseudogenes; the CYP4 clan had 27 intact genes, four pseudogenes, and two partial genes; and the mitochondrial CYP clan had 10 intact genes (Table 1). Across all CYPs, 66% were in genomic clusters (Figure 3). Looking at the four major clans separately, the percentage of clustered genes were: 33% for CYP2, 81% for CYP3, 55% for CYP4, and 50% for mitochondrial CYP.

The CYP gene family had five *Neodiprion*-specific clades with at least five genes (Figure S4a,b), four of which were in the CYP3 clan. Of these, two clades that were both within the CYP3 clan (CYP clades 3 and 5) had evidence of branch-specific, but not site-specific, positive selection (Table 2). CYP clade 3 contained members of the CYP6 subfamily, and the CYP clade 5 contained members of the CYP336 subfamily. Several studies to date suggest that members of the CYP3 clan—and the CYP6 subfamily in particular—play an important role in detoxifying pesticides and host-plant allelochemicals (Feyereisen 2012).

Orthologs were found for all the Halloween genes (which include genes from both the CYP2 and mitochondrial CYP clans) of the 20-hydroxy ecdysone biosynthesis pathway: *CYP302A1* (disembodied), *CYP306A1* (phantom), *CYP307A2* (spookier), *CYP307B1* (spookiest), *CYP314A1* (shade), *CYP315A1* (shadow), and *CYP18A1* which turns over 20-hydroxy ecdysone (Rewitz et al. 2007; Feyereisen 2011; Guittard et al. 2011; Qu et al. 2015). The juvenile hormone biosynthesis gene *CYP15A1* was present as well (Helvig et al. 2004). Finally, *N. lecontei* had orthologs for the two CYP4G enzymes that synthesize the cuticular hydrocarbons used as external waterproof coating (Qiu et al. 2012).

#### Immunity

Antimicrobial peptides (AMPs) are expressed upon infection to kill or inhibit microbes. Based on hymenopteran sequences, the *N. lecontei* AMP gene family had 21 genes (Table 1; Table S6), including single copies of *Hymenoptaecin*, *Abaecin*, and *Tachystatin*, but no clear *Defensin* ortholog. Over 18 *Hisnavicin* genes were identified, including a *Neodiprion*-specific expansion of eight histidine-rich paralogs orthologous to *Hisnavicin-4*, which has been characterized as a larval cuticle protein and AMP, but not functionally tested (Tian et al. 2010). The *N. lecontei* Hisnavicins had a conserved 62 amino acid motif that appeared up to 19 times in a single protein; the purpose of this amplification is unknown. 95% of the AMPs were in genomic clusters (Figure 3). Due to low bootstrap support on many of the branches in our Hisnavicin protein tree, we could not identify unambiguous *Neodiprion*-specific clades (Figure S5a). However, our *Neodiprion* cDNA tree (Figure S5b) did reveal strong support for the monophyly of a cluster of 15 *Hisnavicins* on linkage group 5 (Figure 3), and this cluster had some evidence of positive selection (Table 2).

Outside of the AMP family, most immune pathways had direct orthologs between *N. lecontei* and *D. melanogaster* (Figure S6, Table S7). The basic viral siRNA response pathway was completely conserved between species. The immune deficiency (IMD) pathway was missing an ortholog for the peptidoglycan recognition receptor *PGRP-LC*, but it is likely that another *PGRP* replaced *PGRP-LC* in *N. lecontei*; assigning PGRP orthology was also difficult in ants (Gupta et al. 2015). Also missing is the *Drosophila* mitogen activated protein kinase kinase kinase, TGF-β activated kinase 1 (*Tak1*), but *N. lecontei* had a similar TGF-β activated kinase that is a close ortholog to several *Tak1-like D. melanogaster* proteins possibly involved in immune deficiency signaling. The encapsulation/melanization pathway was missing one of the two *Drosophila* GTPases (*Rak2*). The *N. lecontei Rak1* ortholog may be playing both roles, but again this is likely due to the difficulty of assigning one-to-one orthologs. The Duox pathway was missing the top G-protein coupled receptor, but this is unknown in *D. melanogaster* and unidentified in other Hymenoptera (Evans et al. 2006). Interestingly, *N. lecontei* had two copies of Dual Oxidase (*Duox*), which regulates commensal gut microbiota and infectious microbes (Ha et al. 2005; Lee et al. 2015); *Apis mellifera* had one copy. Finally, the Toll pathway *NF-kappaB* transcription factor, *Dorsal-related immunity factor (Dif)* does not have a one-to-one ortholog in *N. lecontei*, but two copies of its paralog, *Dorsal*, were present.

### Within-genome signatures of adaptive expansions and contractions

#### Evidence of selection in Neodiprion-specific gene family clades

Massive gene family expansions with dozens of genes were not found in *N. lecontei* (in contrast to (Smadja et al. 2009; Zhou et al. 2015)). Instead, the largest *Neodiprion*-specific clade had 22 genes (CYP gene family) and the rest had fewer than 10 genes. Nevertheless, we did identify 11 *Neodiprion*-specific clades containing at least 5 closely related paralogs and a monophyletic clade of 15 AMPs with ambiguous ancestry (Table 1). Of these 12 clades, four had significant branch positive selection (OR clade 1, GR clade 3, CYP clade 3, and CYP clade 5) (Table 2). Of these four clades, only one also had significant site-specific positive selection (GR clade 3) (Table 2).

#### Clustering

Our five focal gene families varied in the proportion of genes that were found in clusters of two or more genes (Fisher’s exact test, *P* = 0.002; Table 1). Post-hoc tests revealed that much of this variation was due to differences between the highly clustered AMP family and all other families except GR (AMP vs. OR: *P* = 0.0091; AMP vs. OBP: *P* = 0.0053; AMP vs. CYP: *P* = 0.024, AMP vs. GR: *P* = 0.12; all p-values are FDR-corrected). The only other difference in clustering that we detected was between the GR and singleton-heavy OBP families (FDR-corrected *P* = 0.045).

Differences in clustering were even more pronounced when we separated the CYP family by clan (Fisher’s exact test, *P* < 0.0001; Table 1). In addition to the pairwise differences described above, we also found that the proportion of CYP3 genes found in clusters differed significantly from ORs, CYP2s, and CYP4s (all FDR-corrected *P* < 0.05), but not AMPs, GRs, and mitochondrial CYPs. Additionally, AMP clustering differed from CYP2, CYP4, and mitochondrial CYP, while GR differed from CYP2 (all FDR-corrected *P* <0.05). Together, these analyses identified AMP, GR, and CYP3 as having an unusually high proportion of genes found in clusters compared to other environmentally responsive gene families.

#### Pseudogenization

Overall, we found very few pseudogenes, and the proportion of pseudogenized genes did not differ significantly among gene families (Fisher’s exact test, *P* = 0.12; Table 1). The chemoreceptors had one pseudogene each while CYP had 12, which is about 10% of the family, but this was also the largest gene family. Although CYP3 had more pseudogenes than other CYP clans, the proportion of pseudogenized genes still did not differ when we compared CYP clans (Fisher’s exact test, *P* = 0.10). Given these low rates of pseudogenization, it is unlikely that *N. lecontei* gene families underwent substantial, recent contractions.

### Ecological correlates of gene-family size across insects

We first examined broad-scale variation in the sizes of our five focal gene families and four CYP clans among different insect orders (Figure S7). Not surprisingly, sample sizes were highly variable across gene families and insect orders. Despite this variation, we observed some intriguing differences among gene families and taxa. We detected significant differences in gene family size among orders for OR (Kruskal-Wallas chisq = 48.2, df = 12, *P* < 1 x 10^−5^), GR (K-W chisq = 25.5, df = 9, *P* = 0.0025), and OBP (K-W chisq = 37.6, df = 9, *P* < 1 x 10^−4^), but not CYP (K-W chisq = 10.3, df = 7, *P* = 0.17) or AMP (K-W chisq = 7.93, df = 5, *P* = 0.16). We note, however, that the AMP sample size was considerably smaller than the other gene families. When we looked at CYP clans individually, we found differences among orders for CYP4 (K-W chisq = 19.0, df = 7, *P* = 0.0083) and mitochondrial CYP (K-W chisq = 16.3, df =7, *P* = 0.022), but not CYP2 (K-W chisq = 9.19, df = 7, *P* = 0.24) or CYP3 (K-W chisq = 8.76, df = 7, *P* = 0.27).

For the OR family, among-group differences in gene number were mostly attributable to an unusually large number of OR genes in Hymenoptera (significant post-hoc tests include Diptera vs. Hymenoptera: *P* = 0.0018; Hemiptera vs. Hymenoptera: *P* = 0.00014; and Odonata vs. Hymenoptera: *P* = 0.011; all p-values are FDR-corrected). By contrast, the size of the OBP family was larger in Diptera than other orders (significant post-hoc tests include Diptera vs. Hymenoptera: *P* = 0.00037; Diptera vs. Hemiptera: *P* = 0.00092; all p-values are FDR-corrected). Although none of the post-hoc tests were significant for GR family size, the Blattodea appear to have more GRs on average than other insect orders (Figure S7). For CYP clans, posthoc tests revealed that hymenopterans have fewer CYP4s than dipterans (FDR-corrected *P* = 0.010) and fewer mitochondrial CYPs than both dipterans and lepidopterans (FDR-corrected *P* = 0.024 and 0.023, respectively).

We next examined how gene family size correlated with ecology within the hymenopteran clade (Figures 4 and 5). Once again, we observed differences among gene families. We found that the number of ORs differed significantly among hymenopteran species that differed in diet (Kruskal-Wallas chisq = 15.8, df = 3, *P* = 0.0012) and sociality (Wilcoxon rank-sum test W = 115; *P* = 0.00094). For diet, we found that herbivores had fewer ORs than all other diet types (fungivores vs. herbivores: *P* = 0.015; omnivores vs. herbivores: *P* = 0.015; insectivores vs. herbivores: *P* = 0.048; all p-values are FDR-corrected). We observed an even more striking difference between eusocial and non-eusocial hymenopterans, with the former having larger OR families, on average. By contrast, GR family size was related to diet (Kruskal-Wallas chisq = 11.8, df = 3, *P* = 0.0082), but not sociality (Wilcoxon rank sum test W = 30; *P* = 0.65). And CYP family size was related to sociality (W = 2; *P* = 0.045), but not diet (*P* = 0.38). Finally, specialists and generalists did not differ significantly in gene family size in any of the gene families and ecology was unrelated to gene family size for OBP and CYP (total CYP number and individual CYP clans). Although these analyses have several limitations (see discussion), these results are consistent with the hypothesis that environmentally responsive gene families may contract or expand predictably in response to particular selection pressures.

**Figure 4.**
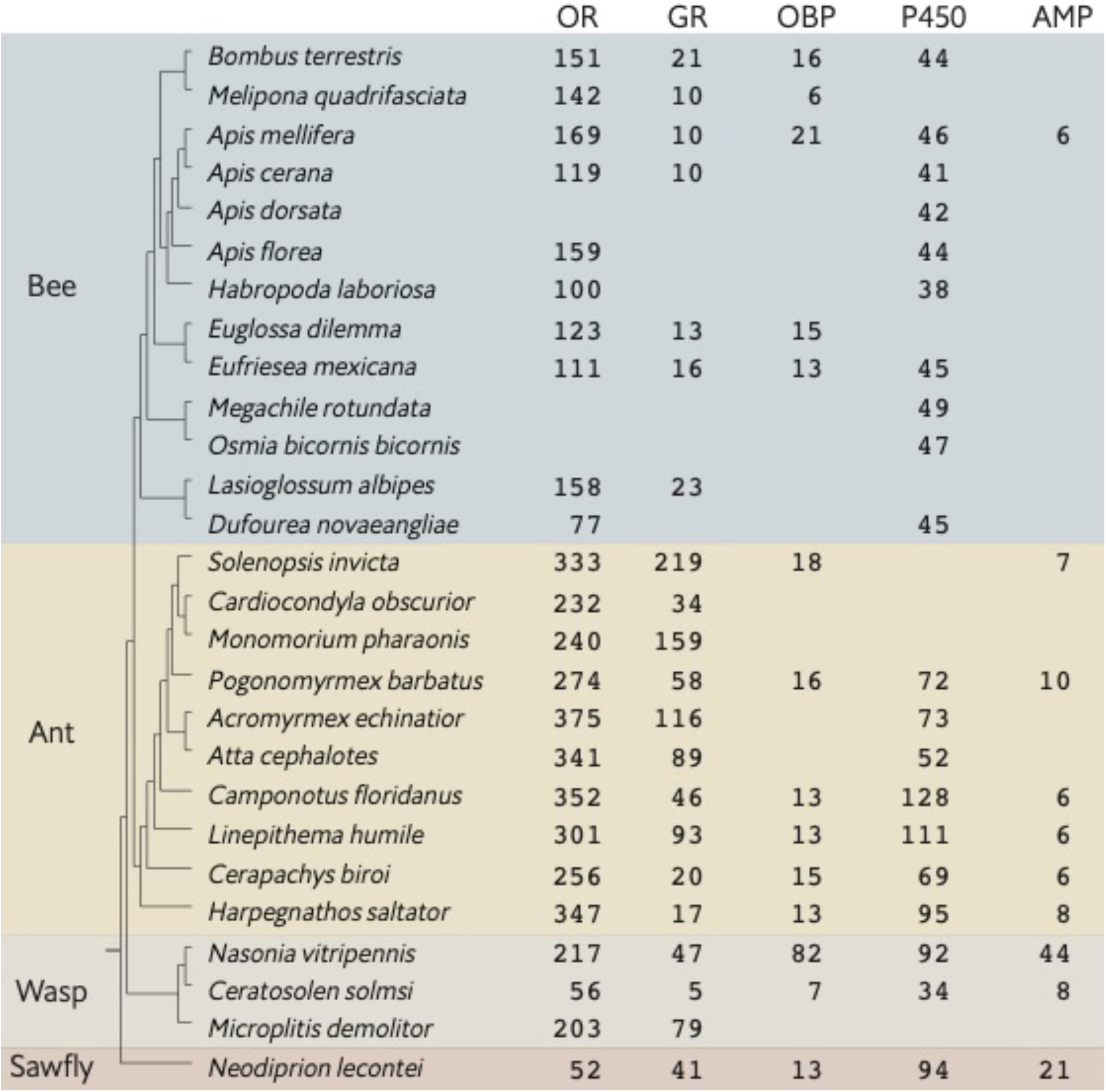
Number of intact genes in hymenopteran genomes for each of five environmentally responsive gene families. Phylogenetic relationships are as in Moreau et al. (2006); Hedtke et al. (2013); Roux et al. (2014); Brand et al. (2017); Branstetter et al. (2017); Peters et al. (2017). Branch lengths are arbitrary. Gene family abbreviations are as in Figure 3.

**Figure 5.**
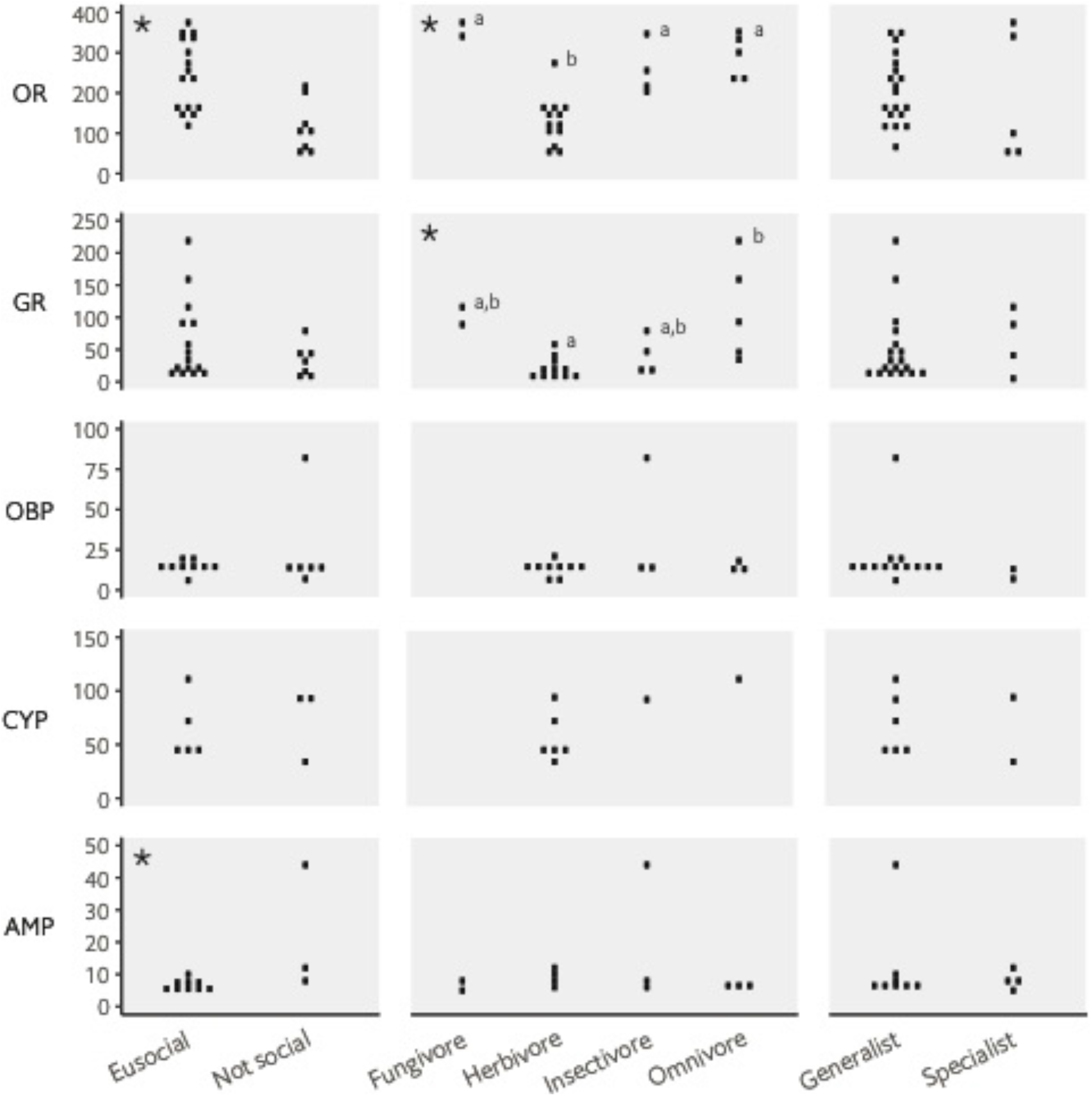
Ecological correlates of gene family size in Hymenoptera. Each point represents the number of intact genes for a hymenopteran species for which both manually curated gene annotations and ecological data are available. Asterisks indicate that gene number varies significantly among the ecological categories under consideration; for significant categories with >2 groups, letters indicate significance in post-hoc tests (groups that do not share a letter are significantly different). Gene number and ecological data for all taxa are provided in Table S8.

## Discussion

The predictability of gene family expansion or contraction in response to specific selection pressures is still an open question. Here, we evaluated genomic signatures of adaptive gene family size changes in five environmentally responsive gene families within the *N. lecontei* draft genome, a hymenopteran exemplar of a pine-specialized lineage. Although we saw minimal evidence of recent gene loss via pseudogenization, at least three gene families (AMP, GR, and CYP3) had genomic distributions consistent with the selective maintenance of novel gene duplicates, and two of these families also had evidence of positive selection within *Neodiprion*-specific clades (GR and CYP3). Next, we examined these same gene families in other hymenopterans to see if family size correlated with diet, ecological specialization, or eusocial behavior. Among Hymenoptera, we found that OR family size was correlated with eusociality and diet type, but not dietary specialization; GR family size was correlated with diet type; and AMP family size was associated with eusociality. These results suggest that ecology can have a predictable impact on gene family size and that different selection pressures impact different gene families. Below, we discuss both the implications and limitations of our analyses and suggest priorities for future comparative work on gene family size evolution.

### Within-genome signatures of gene-family size change

During a niche shift, new selective pressures can leave footprints in the genomes of evolving lineages; such signatures of positive selection are well described for individual loci (Nielsen et al. 2005; Vitti et al. 2013). Similarly, strong selection for increases or decreases in the size of a particular gene family should also leave characteristic genomic footprints. We argue that these footprints include monophyletic groups of closely related paralogs in gene-family trees (from the selective maintenance of novel duplicates), genomic clustering (when novel genes arise via unequal crossing over), evidence of positive selection among paralogs (given selection for sub- or neofunctionalization), and high rates of pseudogenization (from the selective maintenance of loss-of-function mutations). Of the environmentally responsive gene families we evaluated, none exhibited patterns consistent with selection for a decrease in gene family size. By contrast, at least three families had characteristics consistent with selection for an increase in gene family size. Two of these families, GR and CYP3, were highly clustered in the genome and exhibited evidence of positive selection, making these especially promising candidates for expansions related to a novel coniferous host. Additionally, although the AMP family lacked evidence of positive selection, its unusually clustered distribution in the *Neodiprion* genome could be related to selection for increased dosage of a conserved protein function (Perry et al. 2007). Below we discuss the functions of these three candidate families in more detail.

Shifts to pine feeding likely involved changes in the detection of and response to pine-specific cues. Intriguingly, the one GR clade with evidence of positive selection—GR clade 3—is an expansion of six paralogs (one is pseudogenized) orthologous to *DmGR66a*, a bitter receptor specifically for caffeine (Moon et al. 2006). However, orthologs were not found for *DmGR93a* (Lee et al. 2009) and *DmGr33a* (Moon et al. 2009), coreceptors possibly required for caffeine detection. Nevertheless, honeybees, which also lack clear orthologs to these putative coreceptors (Wanner and Robertson 2008), can detect and even prefer low concentrations of caffeine and nicotine (Singaravelan et al. 2005, but see de Brito Sanchez 2011). Although pines do not contain caffeine, they do synthesize alkaloids that could confer some bitterness (Mumm and Hilker 2006). Thus, despite lacking caffeine coreceptor orthologs, members of GR clade 3 may still be involved in the detection of pine-specific bitter compounds. Duplications of putative bitter GRs are documented in other host-specialized insects, such as *Heliconius, Danaus,* and *Bombyx* butterflies (Wanner and Robertson 2008; Briscoe et al. 2013). Our sawfly-specific GR expansion, coupled with the finding that GR family size is associated with diet (see below), lends support to the hypothesis that expansions of GR bitter receptors repeatedly contribute to changes in oviposition and feeding behaviors in plant-feeding insects.

Because pines contain toxic components like terpenoids and phenolics, detoxifying gene families are also promising candidates for pine adaptation. The mountain pine beetle (*Dendroctonus ponderosae*), feeds on pine bark and wood and has gene “blooms” (species-specific gene gains) in the CYP3 and CYP4 clans (Keeling et al. 2013). Similarly, in *N. lecontei*, the CYP family had five blooms (Figure S4a): four CYP3 and one CYP4. CYP3 blooms are also found in wood-feeding insects that do not use pine, such as the emerald ash borer (*Agrilus planipennis*) (David Nelson, unpublished data) and the Asian longhorned beetle (*Anoplophora glabripennis*) (McKenna et al. 2016). Notably, *N. lecontei* larvae frequently ingest pine bark in addition to pine needles (Wilson 1992), suggesting that CYP3 may expand predictably in wood feeders. Additionally, one of the two *Neodiprion*-specific CYP3 clades with evidence of positive selection (Table 3) belongs to the CYP6 subfamily, which is linked to host plant adaptation in several insect taxa (Li et al. 2003; Li et al. 2007; Feyereisen 2012; Mittapelly et al. 2019).

Because pine resin has antimicrobial (Himejima et al. 1992; Cowan 1999; Gershenzon and Dudareva 2007) and fungicidal properties (Grayer and Harborne 1994), we hypothesized that *N. lecontei* co-opted these compounds for its own defense, leading to relaxed selection on genes involved in immunity and a reduced innate immune response. In other Hymenoptera, honeybees (*Apis mellifera*) exposed to plant resin have reduced expression of immune-related genes (Simone et al. 2009) and wood ants (*Formica paralugubris*) that use conifer resin as building material have slightly reduced inducible immune system activity and nests with lower bacterial and fungal loads (Castella et al. 2008). In Diptera, AMP loss is associated with herbivorous lineages that live within host tissue, a more sterile habitat than is experienced by most dipterans (Hanson et al. 2019). Unexpectedly, we found a large species-specific clade of *Hisnavicin*-like AMPs in *Neodiprion*. Although additional data are needed to confirm that *Hisnavicin* orthologs act as AMPs in *N. lecontei*, one possible explanation for this putatively adaptive expansion that lacked an accompanying change in non-synonymous substitution rate is that having large numbers of *Hisnavicin*-like AMPs confers protection against pathogens unique to pine trees. That said, our data do not rule out adaptive AMP loss. For example, *N. lecontei* lacks a clear *Defensin* ortholog, a gene present in all dipterans tested to date (Hanson et al. 2019).

#### Limitations of within-genome analyses

One benefit to studying adaptive expansions/contractions within a single taxon is that gene families have likely experienced similar demographic histories, which can also impact gene birth and death rates. That said, each of our within-genome signatures of selection has limitations that should be revisited with additional data. First, our analysis of genomic clustering does not account for local recombination rate variation, which correlates with tandem array size in several taxa (Gaut et al. 2007). A fine-scale recombination rate map, coupled with clustering analyses for many additional gene families, would more rigorously test the extent to which individual gene family clustering deviates from the genome-wide relationship between recombination rate and tandem array size.

Second, a lack of comparable data from other Eusymphyta meant that our gene family phylogenies lacked orthologues from closely related sawfly taxa. Thus, the “*Neodiprion*-specific” clades may not be unique to pine-feeding sawflies. If these paralogs were present prior to the shift to pine hosts, this would not support a scenario in which new duplicates were selectively maintained in the novel niche. Signatures of positive selection may still be related to pine adaptation but would indicate selection on preexisting loci rather than selection favoring gene family expansion.

Third, signatures of adaptive gene family expansions and contractions may be ephemeral, and the shift to pine use could have occurred too long ago to detect these signatures in *N. lecontei*. For example, in *Drosophila*, pseudogenes have an estimated half-life of ~14.3 million years (Petrov et al. 1996; Petrov and Hartl 1997, 1998). If the rate of gene decay is similar in *Neodiprion*, then pseudogenes that formed after a shift to pine (up to 60 mya) may no longer be detectable in the genomes of extant sawflies. Likewise, gene clustering patterns are likely to change over time from chromosomal rearrangements and additional gene duplications and deletions. To investigate how the number and position of genes in these focal families has changed over time, high quality gene annotations for diprionids and many additional sawfly outgroups are needed. Fortunately, even if footprints of recent gene family size changes are too ephemeral to be detected in most taxa, consistent relationships with ecology should still be detectable given sufficient sampling of taxa differing in ecological traits of interest.

### Ecological correlates of gene family size among hymenopteran taxa

The largest insect OR gene families are in eusocial Hymenoptera, leading to the hypothesis that OR family size expansions were favored in these lineages because they facilitate complex chemical communication (Robertson and Wanner 2006; LeBoeuf et al. 2013; Zhou et al. 2015). To date, evidence in support of this hypothesis has been mixed (e.g., (Roux et al. 2014; Brand and Ramírez 2017). Consistent with the OR-eusociality hypothesis, we found that, on average, eusocial hymenopterans had larger OR families than non-eusocial hymenopterans. However, it is likely that eusocial taxa differ from non-eusocial taxa in many other aspects of their ecology that should also impact OR evolution. Indeed, we found that herbivorous hymenopterans tended to have fewer OR genes than non-herbivores.

Whereas all eusocial hymenopterans had relatively large OR families, some eusocial hymenopterans had relatively small GR families (Figures 4, 5; (Zhou et al. 2015)). To explain the strikingly small set of GR genes in honeybee, Wanner and Robinson (2006) proposed that a stable hive environment and a mutualistic relationship with flowering plants resulted in a lack of selection for GR expansions. Intriguingly, our data indicate that among hymenopterans, GR family size is associated with diet, but not eusociality. Like the ORs, GR gene family size tends to be smaller in herbivores than in non-herbivorous taxa, regardless of social behavior. The directionality of this change, however, is unclear: do shifts to plant diets favor reductions in GR families, do shifts to non-plant diets favor GR expansions, or is it both? Answering this question will require characterizing GR families across many independent transitions to and from herbivory, as well as polarizing directions of change (i.e., distinguishing GR gains from GR losses). Fortunately, there are many such diet transitions across diverse clades of insects (Wiens et al. 2015).

Unlike sociality and diet, ecological specialization was not associated with gene family size in any of the five gene families we evaluated. This result was unexpected because specialization-associated reductions in gene family size are documented in diverse taxa and multiple gene families, including the families examined here (McBride 2007; Smadja et al. 2009; Cao et al. 2014; Goldman-Huertas et al. 2015; Suzuki et al. 2018). One explanation for the lack of association between gene family sizes and specialization in our data is that our “generalist” and “specialist” categories are not meaningful across diverse diets (Forister et al. 2012). Additionally, within a particular diet, the degree of specialization may be highly labile, with rapid fluctuations that are not captured in our broad, order-wide comparison. Indeed, previous studies that reported correlations between gene family size and ecological specialization focused on closely related species. Thus, to fully understand how changes in ecology shape gene family evolution, it will be necessary to evaluate ecological correlates of gene family size at multiple time scales of taxonomic divergence.

Compared to ORs and GRs, our other focal gene families had far less manual annotation data available for analysis. This may explain, in part, why we did not detect strong ecological correlates for the other gene families. It is also possible that by focusing on the sizes of entire gene families, we missed relevant signals in particular subfamilies (Hahn et al. 2007). For example, as noted above, expansions of CYP3 and CYP4 subfamilies are associated with wood-feeding insects and CYP3 clan subfamilies were also linked to detoxification in honey bee (Berenbaum and Johnson 2015; Johnson et al. 2018). However, we did not detect any correlations between ecology and CYP clan sizes. Despite these limitations, we did uncover hints that AMP gene family size may be larger in non-eusocial lineages. If eusocial taxa tend to inhabit more sterile environments (nests and hives) than non-eusocial taxa, this finding is consistent with associations between habitat and AMP loss reported in dipterans (Hanson et al. 2019). Given that AMPs were also implicated in our within-genome analysis, immune-related genes are especially promising candidates for future manual annotation projects.

#### Limitations of among-taxon analyses

Comparative analysis is a powerful approach for evaluating the repeatability and predictability of evolutionary outcomes. Although our comparison of candidate gene family sizes among ecologically diverse hymenopterans hints at intriguing relationships between ecology and gene family size, it also had several limitations that should be revisited in future work. First, because several taxa in our manual annotation dataset are missing from published hymenopteran phylogenies (Peters et al. 2017), we were unable to correct for phylogenetic non-independence and polarize gene gain/loss (e.g., as in (Hahn et al. 2005; Han et al. 2013) without losing unacceptable amounts of data. Without accounting for similarity in ecology and gene family size due to recent common ancestry, our Type I error rate is likely inflated and p-values should be interpreted with caution. Nevertheless, variation in patterns of association among ecological traits and gene families suggest that phylogeny and ecology are, to some extent, decoupled.

The gene annotation and ecological datasets also had limitations. For example, across studies that included manual annotations, we observed a lack of consistency in the methods and criteria for manually curated gene family datasets. The most problematic inconsistency was in the criteria for delineating intact, partial, and pseudogenized gene annotations. “Intact” could mean an exon-by-exon check against closely related orthologs, a minimum amino acid length, or merely the presence of an expected domain. Meanwhile, in reference publications, the number of pseudogenized and partial annotations were not always reported or were conflated. This is in addition to variation in the methods used to search for genes. Inconsistency in annotation methods and criteria across studies may introduce taxon-specific biases unrelated to ecology. Regarding ecology, categorizations are somewhat subjective. For example, this study and Rane et al. (2016) classified bees as generalists since they collect nectar and pollen from multiple plant families (we defined specialization as the use of a single taxonomic family). But Johnson et al. (2018) classified bees as specialists as their diet consists of only nectar and pollen.

Finally, our attempts to correlate the size of different gene families with ecology suffered from sampling biases in which species had genome assemblies and which gene families were manually annotated. Species skewed heavily towards *Drosophila* and apocritan Hymenoptera, and annotations toward the OR and CYP families (Table S8). To evaluate ecological correlates of gene family expansions and contractions, it is essential to expand both the taxonomic breadth and depth of annotation sampling. Taxa that capture independent ecological transitions (e.g., between herbivory and other diets) would be especially useful, as would replicated groups of closely related species that vary in ecological axes of interest (e.g., specialization or social behavior). By systematically sampling different ranges of divergence times, we can evaluate the extent to which the tempo of gene family size change varies across different gene families. To do so, however, will require high quality, manually curated datasets produced using consistent methods and standards for many different environmentally responsive gene families.

## Conclusions

Gene families that mediate ecological interactions may predictably expand and contract in response to changing selection pressures. These adaptive changes in gene family size should leave detectable genomic footprints in recent niche colonists and across taxa with convergent niche shifts. Consistent with these predictions, (1) our analysis of gene family evolution in a derived pine feeder suggests that expansions of GRs, CYP3s, and AMPs may have accompanied pine adaptation, and (2) our comparison among ecologically diverse hymenopterans links two of these families to variation in diet (GR) and eusociality (AMP). In the order Hymenoptera, the OR gene family was associated with ecology (eusociality), however, the size of all five candidate gene families was not linked to other ecological axes of variation (specialization/generalization); they were in other comparisons of closely related species (McBride 2007)). Together, these results suggest that the size changes of environmentally responsive gene families vary in both temporal dynamics (shallow vs. deep divergence times) and in ecological drivers. Teasing apart these relationships will require high quality annotation data across diverse gene families, ecologies, and divergence times. For hymenopterans, increased effort in understudied symphytan, parasitoid, and herbivorous taxa would be especially useful for disentangling different axes of ecological variation contributing to gene family size change.

## Materials and methods

### Biological material

To minimize the confounding effects heterozygosity has on genome assembly, we sequenced haploid siblings. Like all Hymenoptera, sawflies have haplodiploid sex determination in which males (haploid genomes) emerge from unfertilized eggs and females (diploid genomes) from fertilized eggs. A virgin female will bear a clutch of all-male offspring with haploid recombinants of the maternal genome. But the individual genomes are not identical, so an assembly derived from a single clutch is akin to a diploid assembly made from a single individual.

All insects were reared in custom, climate-controlled environmental chambers (18:6 light cycle, 22°C, 70% RH) on jack pine (*Pinus banksiana*) foliage. Our laboratory line of *N. lecontei* was established from multiple larval colonies collected from a mugo pine (*P. mugo*) in Lexington, Kentucky, USA (37°59’01.6”N 84°30’38.8”W; population ID: RB017). For the transcriptome, adults and larvae were collected from the first lab-reared generation; both were stored at −80°C. For the genome assembly, the founding population was propagated in the lab for two generations, followed by brother-sister matings for an additional two generations. At this point, a single, virgin, adult female (I2G2-V, 4^th^ generation in the lab) was allowed to lay unfertilized eggs onto jack pine seedlings. The offspring (haploid male brothers from an inbred mother) were reared until the eonymph (prepupal) life stage, at which point they were isolated without food for 24 hours prior to preservation in absolute ethanol at −20°C. Although eonymphs are non-feeding, they were starved to ensure the gut contents were completely voided.

### Sample preparation and sequencing

#### Genomic DNA

Whole eonymph bodies were individually frozen inside microcentrifuge tubes with liquid nitrogen and ground with pestles made from 1-mL micropipette tips; the resulting powder was incubated in CTAB buffer supplemented with proteinase K and RNase A. After PCI extraction and ethanol precipitation, the precipitate was dried overnight before being resuspended in TE buffer. DNA integrity was assessed with 0.7% agarose gel, purity was measured with the 260/280 ratio, and concentration was measured with a Quant-iT dsDNA High-Sensitivity fluorescence assay (Thermo Fisher Scientific).

The HudsonAlpha Genomic Services Lab (Huntsville, AL, USA) prepared and sequenced the DNA libraries. Two small-insert, barcoded libraries with average fragment sizes of 337 bp and 864 bp were made from a single individual. A 4.6-kbp mate-pair, barcoded library was made from 25 pooled individuals. All individuals were brothers from the same I2G2-V mother. The libraries were sequenced on Illumina HiSeq 2000 with paired-end, 100 bp (PE100) reads: the small-insert libraries each had ¼ of a flow cell lane and the mate-pair library had an entire lane.

#### mRNA

The RNeasy Mini extraction kit (Qiagen) was used to collect total RNA from adult female body, adult female head, adult male body, adult male head, eonymph body, feeding larval body, and feeding larval head. RNA from eonymph head was extracted but not sequenced due to insufficient yield. Each tissue was represented with one replicate that had equal RNA contributions from eight individuals, except for eonymph body which was comprised of three individuals. RNA integrity and concentration were measured with a 2100 Bioanalyzer (Agilent).

The HudsonAlpha Genomic Services Lab (Huntsville, AL, USA) handled library preparation and sequencing. Non-stranded, barcoded libraries were made for each of the seven tissue samples; on average, mRNA was sheared to 200 bp. The libraries were combined and sequenced on an entire flow cell of Illumina MiSeq with PE250 reads in addition to one lane of Illumina HiSeq 2000 with PE50 reads.

### Read processing and assembly

#### De novo genome assembly

Sequencing reads were chastity-filtered and adaptor-trimmed with fastq-mcf (ea-utils v1.04.803) (Aronesty 2011), and quality-filtered with fastq_quality_filter (FASTX Toolkit v0.0.13.2) (Gordon and Hannon 2019). The 337-bp small-insert reads and the 4.6-kbp mate-pair reads were quality-filtered to retain reads where at least 80% of the bases had a quality score of 20 or higher (parameters: **–q 20 –p 80**). Due to sequencing quality, the 864-bp small-insert reads were filtered to retain reads where at least 70% of the bases had a quality score of 20 or higher (R1) or 60% (R2) (parameters: **–q 20 –p 60/70**). In situations where only one end of the paired-end reads passed filtering, the passed reads were kept and treated as single-end data. Kmer counting was used to measure read depth before and after filtering (Jellyfish v1.1.11) (Marçais and Kingsford 2011). Finally, reads were screened for sequencing contamination by mapping the reads (BWA v0.7.12-r1039) (Li and Durbin 2009) to reference genomes for *Escherichia coli* (K12 substr. DH10B uid58979), human (v37), loblolly pine (*Pinus taeda*, v0.8), and *Wolbachia* (endosymbiont of Dmel uid57851).

The genome was assembled with ALLPATHS-LG (v47417) (Gnerre et al. 2011) using default settings, including a minimum scaffold size of 1000 bp. The error-correction module was run on the reads prior to assembly. After assembly, GapFiller (v1.11) (Boetzer and Pirovano 2012) was used to help close intra-scaffold gaps. Spurious scaffolds were identified with SOAP.coverage (v2.7.7) (Li et al. 2009): reads were mapped to the assembly scaffolds and scaffolds with a read depth < 15 and nucleotide percentage < 40 were removed. The completeness of the final assembly was measured with CEGMA (v2.5) (Parra et al. 2007) and BUSCO (v1.22) (Simão et al. 2015) benchmarks. BUSCO was run with the arthropoda-25oct16 database (`parameters: –−long`).

#### De novo transcriptome assembly

For both the PE250 MiSeq and the PE50 HiSeq reads, fastq-mcf (ea-utils v.1.04.803) (Aronesty 2011) was used for chastity filtering and Trimmomatic (v0.32) (Bolger et al. 2014) was used to adaptor clip, trim, and quality-filter. The PE250 MiSeq reads were processed with the Trimmomatic parameters `ILLUMINACLIP: 2:15:5, HEADCROP: 10, CROP: 60, MINLEN: 60, AVGQUAL: 25` whereas the PE50 HiSeq reads were processed with `ILLUMINACLIP: 2:15:5, HEADCROP: 15, MINLEN: 35, AVGQUAL: 25`. Because the mRNA libraries had an average insert size of 200 bp, the MiSeq reads required extensive adaptor trimming. Reads were screened for contamination as described in *De novo genome assembly*.

For each tissue, transcriptomes were assembled with Trinity (r2013_08_14) (Grabherr et al. 2011; Haas et al. 2013) using default settings and the `--jaccard_clip` option. Spurious sequences were identified by mapping the sequencing reads to the assembled transcripts with RSEM (v1.2.18) (Li and Dewey 2011); transcripts with either FPKM or TPM values < 1 were removed. After filtering, the transcriptomes were combined, and duplicate sequences were removed.

### Genome size estimation

Flow cytometry was described in (Harper et al. 2016). For this analysis, we used adult males and females from a lab line of *N. lecontei* established from a colony collected in Auburn, GA (33°59’22.4” N, 83°47’44.6”W; population ID: RB027). Briefly, cell nuclei were collected from the heads of 7 individuals (4 female, 3 male) and stained with propidium iodide. Mean fluorescence for each sample was measured with a BD FACSCalibur flow cytometer (BD Biosciences) and compared to two external standards: *Drosophila melanogaster* (adult female heads, 1C = 175 Mbp) and *Gallus gallus domesticus* (CEN singlets from BioSure, Grass Valley, CA, 1C = 1222.5 Mbp). To correct for ploidy differences between haploid males and diploid standards, we multiplied the *N. lecontei* male estimates by 2. To obtain a single size estimate for each N. lecontei sample, we averaged values obtained for the two standards.

### Repeat annotation

The N. lecontei genome assembly was masked with a custom repeat library. A lineage-specific de novo repeat library was made with RepeatModeler (v1.0.7) (Smit and Hubley 2008-2015) and combined with the hymenopteran repetitive element database (Nov. 2013) from Repbase (Jurka et al. 2005). The custom library was used by RepeatMasker (v4.0.3) (Smit et al. 2013-2015) (parameters: `-cutoff 250 -s -pa 15 -gc 40 -a –poly`) to identify and mask repetitive elements in the genome, including low-complexity DNA and simple repeats.

Transposable element (TE) family consensus sequences were identified by rerunning RepeatModeler (Smit and Hubley 2008-2015) on the genome assembly using the “ncbi” search engine. The resulting sequences were provided to RepeatMasker (Smit et al. 2013-2015) as a custom library to locate associated TE copies in the genome (parameters: `-gc 40 -cutoff 250 -gff -gccalc -norna -nolow -no_is –poly`). TE families with at least 10 fragments longer than 100 bp were extracted for further analysis.

The sequencing reads were mapped to a concatenation of the masked genome and the consensus TE sequences (BWA MEM (parameters: `-M`) (Li and Durbin 2009)). Families that had at least 1x the median coverage to the reference genome for at least 80% of their sequence (to support at least one full insertion found by RepeatModeler) and at least 2x the maximum coverage of the reference genome (to support multiple insertions of the family) were extracted with genomeCoverageBed (BEDtools (Quinlan and Hall 2010)). We attempted to identify the consensus sequences with BLASTN and BLASTX (Altschul et al. 1990) searches against a database of repeat elements, but the only hits were to the lineage-specific elements identified by RepeatModeler. Sequences were also filtered for BLAST hits to rRNA or mitochondrial sequences.

We also used dnaPipeTE (Goubert et al. 2015) to identify what proportion of our short reads was composed of repetitive content, we used a random subset of reads corresponding to 1-fold coverage of the genome (331Mb) and took the total for three separate random samplings of reads (parameters: genome size = 331000000 genome coverage = 1 samples number = 3). We then compared this annotation to the RepeatModeler annotation.

### Gene and functional annotation

#### Automated gene annotation

RNA-Seq data for *N. lecontei* was used to generate training models for gene prediction along with utilization of peptide sequences from other species. PASA (r20130425beta) was used to build a comprehensive transcriptome set from Trinity assembled transcripts along with RNA-Seq read mapping predictions generated from the Tuxedo pipeline. To improve annotation quality, in addition to this *N. lecontei* transcriptome, annotated proteins from *Atta cephalotes* (OGSv1.2), *Acromyrmex echinatior* (OGSv3.8), *Apis melifera* (OGSv3.2), *Athalia rosae* (OGSv1.0), and *Nasonia vitripennis* (OGSv1.0) were provided to Maker (2.09) (Cantarel et al. 2008) as evidence for structural gene prediction. Prior to annotation, the genome was masked using a custom repeat database built using RepeatModeler (v1.0.8) and the annotation was run using the *ab initio* gene predictors Augusts, Genemark-ES and snap in addition to the evidence provided. The functions of the predicted protein-coding genes were putatively established with BLASTP alignments (Altschul et al. 1990) to the Swiss-Prot database (accessed 20 Apr 12) (Apweiler et al. 2004). In cases of multiple matches, the top-ranked alignment was assigned to the gene annotation. Protein motifs and functional domains within the annotations were also identified with an InterProScan (v5.3.46.0) (Jones et al. 2014) search against the InterPro database with gene ontology and IPR lookup (Finn et al. 2016). For the official gene set (OGS), the Maker annotations were filtered by hits to the reference databases and/or a minimum eAED score of 0.1. A second set of gene annotations was generated with the NCBI GNOMON pipeline (annotation release 100 on Nlec1.0 assembly, GCF_001263575.1) (Souvorov et al. 2010).

As the genome was annotated prior to submission to NCBI, we encountered a problem when the NCBI contamination software flagged vector/adaptor sequences for removal; this would disrupt the coordinates provided by Maker. We used a modified version of GAG (Hall et al. 2014) that could accept the flagged coordinates from NCBI to edit the assembly and update annotation coordinates accordingly.

#### Chemoreceptor genes

The olfactory (OR) and gustatory (GR) receptor genes were manually curated following Robertson et al. (2003, 2006). Amino acid sequences of manually curated chemoreceptor genes from *Apis mellifera* (Robertson and Wanner 2006; Smith et al. 2011), *Bombus terrestris* (Sadd et al. 2015) and *Cephus cinctus* (Robertson et al. 2018), *Drosophila melanogaster* (Flybase release FB2017_04), and *Nasonia vitripennis* (Robertson et al. 2010) were used as queries in TBLASTN (v2.2.19) (Altschul et al. 1990) searches against the *N. lecontei* draft genome (parameters: `-e 100000 -F F`). Gene models were manually built in TextWrangler (v5.5) (Bare Bones Software), using protein alignment to identify exons and refine the gene structures; alignments were visualized with Clustal X (v2.1) (Larkin et al. 2007). The Neural Network Splice Predictor program from the Berkeley *Drosophila* Genome Project was used to help identify intron splice sites (http://www.fruitfly.org/seq_tools/splice.html). New gene models were added to TBLASTN searches and this process continued iteratively until new chemoreceptors were no longer found. The gene models were checked against RNAseq reads from tissue-specific transcriptomes (adult antennae, mouthparts, heads, legs, genitalia, and larval heads (Herrig et al. 2019)) and against orthologs in the *N. pinetum* draft genome assembly (NCBI accession GCA_004916985.1).

#### Odorant binding proteins

Custom scripts were used to identify Maker gene annotations (see *Automated gene annotation*) that contained the classic/6C, Plus-C, Minus-C, or atypical odorant binding protein (OBP) motif (Xu et al. 2009). These as well as OBPs from *Apis mellifera* and *Nasonia vitripennis* were used as queries for TBLASTN searches against the *N. lecontei* genome; searches did not yield any new OBPs. All genomic regions identified as potential OBPs were manually curated as described for chemoreceptor genes. After manual annotation, duplicate annotations or genes that lacked OBP motifs were removed.

#### Cytochrome P450 genes

A broad set of 52 insect CYP genes (covering the diversity of insect CYP families) were searched against the *N. lecontei* genome assembly (E-value cutoff 1e3). Scaffolds with hits were then searched against 8782 known insect CYPs. The top 10 hits were returned (later increased to 15 to recover more sequences) and filtered for duplicates. An alternative search of the NCBI GNOMON predictions (“Neodiprion lecontei[orgn] AND P450 NOT reductase”) was also performed and new sequences were added to the dataset. This approach found all the loci identified by the initial search, indicating that the GNOMON annotation tool was able to comprehensively search for CYP sequences. Finally, the candidate *N. lecontei* CYP sequences were manually curated based on comparison to the best BLAST hits.

#### Immune-related genes

Because of the relative completeness of its immune annotation, *Drosophila melanogaster* immunity genes were used to guide annotation. Reference immune genes from *D. melanogaster* tagged with the gene ontology term “GO:0002376 – Immune system process” were compiled from Flybase (release 6.13). Orthology with *N. lecontei* proteins was assigned initially with reciprocal BLASTP (Altschul et al. 1990) searches (E-value cutoff 1e-10). Reference *D. melanogaster* genes without obvious one-to-one orthologs in *N. lecontei* were examined individually to determine whether closely related paralogs in one or both species interfered with the inference of orthology. If not, they were searched against the *N. lecontei* genome assembly using TBLASTN (Altschul et al. 1990) in an attempt to identify unannotated orthologs.

Since antimicrobial peptides (AMP) are unlikely to be conserved between *D. melanogaster* and *N. lecontei*, AMPs from three representative hymenopterans *Apis mellifera* (Danihlík et al. 2015), *Nasonia vitripennis* (Tian et al. 2010), and *Camponotus floridanus* (Ratzka et al. 2012; Zhang and Zhu 2012; Gupta et al. 2015) were used for BLAST queries. Furthermore, since AMP copy number is fast evolving, we attempted to find all the *N. lecontei* orthologs of each hymenopteran AMP instead of focusing on one-to-one orthology. Once again, BLASTP searches were performed against the annotated proteins and TBLASTN searches were performed against the assembled genome; the TBLASTN search did not reveal additional AMPs. Putative *N. lecontei* orthologs were reciprocally blasted against the appropriate hymenopteran proteome to assure that the best hits were indeed AMPs.

Amino acid and cDNA sequences for all manual annotated genes are available in File S1.

### Glomeruli counts

#### Antennal lobe histology

Whole heads of adult *N. lecontei* of both sexes were fixed in 2% paraformaldehyde, 2% glutaraldehyde in PBS for 5 days. Heads were rinsed for 40 minutes three times and the brains dissected out in cold PBS. Following blocking with goat serum, brains were permeabilized with 1% Triton X-100 in PBS (Electron Microscopy Supply, Fort Washington, PA; PBS-TX), rinsed with 0.1% PBS-TX, and incubated on a shaker at 25°C for three nights in primary antibody (1:500 in 2% goat serum in 0.2% PBS-TX). Monoclonal *Drosophila* synapsin I antibody (SYNORF1, AB_2315426) from the Developmental Studies Hybridoma Bank (catalog 3C11) was used to label synapsin. Subsequently, brains were washed in 0.1% PBS-TX and incubated for two nights in Alexa Fluor 568 (ThermoFisher) goat anti-mouse secondary antibody (1:100 in PBS) in the dark at room temperature on a shaker. After secondary incubation, brains were rinsed with distilled water, dehydrated in increasing concentrations of ethanol, and mounted in custom-made aluminum well slides. Brains were cleared by removing ethanol and replacing it with methyl salicylate. Brains were imaged on an inverted Zeiss 880 Laser Scanning Confocal Microscope with a 20X plan-Apochromat 20x 0.8 aperture objective and optically sectioned in the horizontal plane at 3-micron intervals.

#### Glomeruli segmentation

Whole-brain images of one female and one male were manually segmented using the TrakEM2 software package in ImageJ (Cardona et al. 2012; Schindelin et al. 2012). Individual glomeruli were traced in both brain hemispheres. Glomeruli near the center of the antennal lobe can be difficult to distinguish, meaning counts are biased toward fewer glomeruli and the largest number of glomeruli confidently detected represents a minimum of the number of expected glomeruli. Male *Neodiprion* have a collection of smaller synaptic clusters in their antennal lobe (Dacks and Nighorn 2011), but the functional significance of this anatomy is not known. There are more than 50 of these smaller synaptic clusters and we suspect they do not represent the traditional one-to-one OR-to-glomerulus organization. Therefore, these structures were not included in counts. Male glomeruli number may be lower if particular OSNs contribute to these clusters instead of forming traditional glomeruli.

### Within-genome signatures of adaptive expansions and contractions

#### Clustering and pseudogene analyses

To evaluate the extent to which members of our five focal gene families were located in tandem arrays, we placed our annotated genes on a linkage-map anchored version of the *N. lecontei* genome assembly described in Linnen et al. 2018. We considered genes to be clustered if they were located within a genomic region of 20(n - 1) kilobases, where n is the number of genes in the cluster under consideration. This criterion was chosen based on average gene densities in *Nasonia* (Niehuis et al. 2010) and clustering criteria described *Drosophila* (Vieira et al. 2007). For scaffolds that could not be placed on linkage groups, we evaluated clustering only if genes were more than 20 kb from either scaffold end.

To evaluate whether the five focal gene families differed in (1) the proportion of genes found in clusters of two or more or (2) the proportion of pseudogenized genes, we performed Fisher’s exact tests in R v3.5.0 (“fisher.test” function) (R-Core-Team 2018). For significant Fisher’s exact tests, we performed additional posthoc tests using the “fisher.multcomp” function (from R package RVAideMemoire v. 0.9-72) with FDR correction (Benjamini-Hochberg method) for multiple comparisons.

#### *Identification of* Neodiprion-*specific clades and tests of positive selection*

First, we identified clades unique to *N. lecontei*. For each gene family, a multi-species, amino acid phylogeny was constructed with manually curated annotations from *N. lecontei*, select Hymenoptera, and *D. melanogaster*. Sequences were size filtered (350≥ for GR, OR, CYP; 100≥ for histnavicin and OBP), but pseudogenes and partial annotations that met the length requirement were retained. MAFFT alignments (v7.305b) (Katoh et al. 2002) (parameters: `--maxiterate 1000 –localpair`) were visually inspected to remove sequences with large alignment gaps, and sites with more than 20% gaps were removed with trimAl (v1.4.rev15 build[2013-12-17]) (Capella-Gutiérrez et al. 2009) (parameters: `-gapthreshold 0.8`). Maximum likelihood phylogenies were made in RAxML (v8.2.4) (Stamatakis 2014) (parameters: `-f a -x 12345 -p 12345 -# autoMRE`) using protein substitution models chosen from ProtTest3 (v3.4.2) (Abascal et al. 2010; Darriba et al. 2011).

*Neodiprion*-specific clades were defined as those with at least five *N. lecontei* genes (not including partial and pseudogenes) and a bootstrap score ≥70 (Engsontia et al. 2015). Second, the clades were confirmed with cDNA phylogenies for each *N. lecontei* gene family. Amino acid sequences were aligned as above, however, after alignment TranslatorX (Abascal et al. 2010) was used to map cDNA sequences to the amino acid alignment. After trimming, the cDNA alignments were passed to RAxML to construct maximum likelihood gene family trees with the nucleotide substitution model `-m GTRGAMMA`.

Site tests were conducted with codeml (part of the PAML package (PAML v4.9e) (Yang 2007)) using the cDNA phylogenies and sequences as inputs. For each *Neodiprion*-specific clade, the gene family cDNA phylogeny was pruned to remove all branches except for that clade. Codeml models M7, M8, and M8a were fitted to the cDNA sequence and phylogeny data. Likelihood-ratio tests were performed for the nested models M7-M8 (null model M7 that equally distributes amino acid sites across 10 classes of ω parameter values (p, q) against alternative model M8 that has an 11^th^ class for positively selected sites) and M8-M8a (null model M8a that has 11 classes and does not allow positive selection against alternative model M8). Bonferroni correction was applied to the likelihood-ratio test probability values; each value was multiplied by two since two tests that used M8 as the alternative model were performed on each clade. For clades with significant likelihood-ratio tests, sites under selection were identified by looking at the Bayes Empirical Bayes analysis within the alternative models.

For branch tests, the cDNA phylogenies for each *N. lecontei* gene family were used to compare the lineage-specific clade to the rest of the gene family. To determine if the foreground branch dN/dS (i.e., the branch with the species-specific expansion) was significantly different from the background (i.e., the rest of the gene family), in codeml we ran a two-ratio model (Model=2, fix_omega=0) and a one-ratio model (Model=0, fix_omega=0) for that clade and performed a likelihood-ratio test comparing the two models. To determine if the foreground branch is under positive selection (dN/dS>1), we performed a likelihood-ratio test comparing the two-ratio model to a neutral model (fix_omega=1).

### Ecological correlates of gene family size among insects

All the insect genome assembly projects we could find (published and unpublished) were searched for manually curated OR, GR, OBP, CYP, and AMP gene annotations. If fasta sequence files were available, the number of intact, partial, and pseudogenized genes was determined by gene names (e.g., labels with “pse” or “partial”) and compared to values reported in the publication. Otherwise, we relied on reported values. If gene family size was reported but not broken down into intact, partial, and pseudogenized, and sequence files were unavailable, we assumed that the reported number referred to intact genes. Splice variants were not included in the gene count. It is important to note that different authors likely used different criteria for these categories.

Only putatively functional (intact) gene were used in gene family size comparisons. Species were classified according to taxonomic order, diet type, dietary specialization, and sociality. An order needed at least two species to be included. Specialization was defined as the use of a single taxonomic family and only referred to the realized diet niche, ignoring reports of feeding under laboratory conditions. If a species had a preferred host or both specialist and generalist life stages, it was classified as specialist. Comparisons were made in R (v3.5.0) where species were grouped by the different classifications.

Because both gene family size and ecology are likely to correlate with phylogeny, the ideal approach to identifying ecological correlates of gene family evolution is to use statistical methods that account for phylogenetic relationships (Hahn et al. 2005; De Bie et al. 2006; Han et al. 2013). Unfortunately, a lack of overlap between species with manual annotations for our focal gene families and species included in published hymenopteran genomes precluded us from such an analysis without a substantial loss of sample size. Therefore, as a first step to evaluating ecological correlates of gene family size, we used non-parametric tests to determine whether gene family size differed among taxa. For sociality and specialization, we used two-tailed Wilcoxon rank-sum tests (“wilcox.exact” function in the exactRankTests v0.8-30 package). For taxonomic order and diet, both of which have more than two categories, we used Kruskal-Wallis tests (“kruskal.test” function) followed by Dunn’s post-hoc tests of multiple comparisons (“dunnTest” function in the FSA v0.8.23 package).

## Acknowledgements

We thank Linnen lab members for insect collection, insect reading, and reading earlier manuscripts; Jeramiah Smith, Erin Scully, and Romain Studer for advice. We are especially grateful to Hugh Robertson for his guidance on manual chemoreceptor gene annotation. This work was supported by the University of Kentucky Center for Computational Sciences and the Lipscomb High Performance Computing Cluster, the United States Department of Agriculture National Institute of Food and Agriculture (2016-67014-2475; CRL), the Kentucky Science and Engineering Foundation (KSEF-3492-RDE-019; CRL), and the University of Kentucky (Lyman T. Johnson Fellowship; KV).

## Data availability

The genome assembly, official gene set (OGS), and transcriptome described in this paper (v1 versions) can be found at https://i5k.nal.usda.gov/neodiprion-lecontei

On GenBank (NCBI), the genome assembly is labeled whole genome shotgun sequencing project accession PRJNA28045 and the genomic sequencing reads are RefSeq accession PRJNA312506. The transcriptome is transcriptome shotgun assembly accession GEDM00000000; this is a combined transcriptome of all seven tissue types. The mRNA sequencing reads for each tissue type was submitted separately under BioSample and short read archive accessions SAMN04302192 (adult female head), SAMN04302193 (adult female body), SAMN04302194 (adult male head), SAMN04302195 (adult male body), SAMN04302196 (feeding larval head), SAMN04302197 (feeding larval body), and SAMN04302198 (eonymph body). The predicted gene annotations on NCBI are from Gnomon, the NCBI annotation pipeline, and were not described in this paper. Finally, the clustering analysis was based on a linkage-map anchored version of the genome assembly described in Linnen et al. 2018. This anchored assembly is denoted as v1.1 in NCBI and the *N. lecontei* i5k Workspace@NAL (USDA).

