## Supplemental for "Ecological correlates of gene family size in a pine-feeding sawfly genome and across Hymenoptera": Vertacnik_etal_supplementary_data_19Feb21.docx

**Supplementary materials for**

**Investigating the ecological correlates of gene family size evolution using the draft genome of the redheaded pine sawfly *Neodiprion lecontei* (Hymenoptera: Diprionidae)**

Kim L. Vertacnik^1,2^, Danielle K. Herrig^1^, R. Keating Godfrey^3^, Tom Hill^4,5^, Scott M. Geib^6^, Robert L. Unckless^4^, David R. Nelson^7^, and Catherine R. Linnen^1^

^1^Department of Biology, University of Kentucky, Lexington, KY 40506, USA

^2^Columbia River Inter-Tribal Fish Commission, Hagerman, ID 83332, USA (current address)

^3^Department of Neuroscience, University of Arizona, Tucson, AZ 85721, USA

^4^Department of Molecular Biosciences, University of Kansas, Lawrence, Kansas 66045, USA

^5^NIH Frederick National Laboratory for Cancer Research, Frederick, MD 21702, USA

^6^Tropical Crop and Commodity Protection Research Unit, United States Department of Agriculture: Agriculture Research Service Pacific Basin Agricultural Research Center, Hilo, Hawaii 96720, USA

^7^Department of Microbiology, Immunology and Biochemistry, University of Tennessee Health Science Center, Memphis, TN 38163, USA

**Supporting information**

**Supplementary figure S1a:** Olfactory receptor gene family amino acid phylogeny with select Hymenoptera

**Supplementary figure S1b.** Olfactory receptor gene family cDNA phylogeny for *N. lecontei* only

**Supplementary figure S2a.** Gustatory receptor gene family amino acid phylogeny with select Hymenoptera

**Supplementary figure S2b.** Gustatory receptor gene family cDNA phylogeny for *N. lecontei* only

**Supplementary figure S3a.** Odorant binding protein gene family amino acid phylogeny with select Hymenoptera

**Supplementary figure S3b.** Odorant binding protein gene family cDNA phylogeny for *N. lecontei* only

**Supplementary figure S4a.** Cytochrome P450 gene family amino acid phylogeny with select Hymenoptera.

**Supplementary figure S4b.** Cytochrome P450 gene family cDNA phylogeny for *N. lecontei* only

**Supplementary figure S5a.** Hisnavicin gene family amino acid phylogeny with select Hymenoptera

**Supplementary figure S5b.** Hisnavicin gene family cDNA phylogeny for *N. lecontei* only

**Supplementary figure S6.** **An overview of *N. lecontei* immune pathways.**

**Supplementary figure S7. Variation in gene family size among insect orders.**

**Supplementary table S1:** Sequencing libraries and read counts for the *N. lecontei* genome

**Supplementary table S2:** Comparison of genome assembly and annotation statistics from published hymenopteran draft genomes

**Supplementary table S3.** Summary of repetitive elements identified in the *N. lecontei* genome

**Supplementary table S4.** Tissue types, read counts, and transcript counts for the *N. lecontei* transcriptome

**Supplementary table S5.** Glomeruli counts from left and right antennal lobes of an adult *N. lecontei* male and an adult *N. lecontei* female.

**Supplementary table S6.** Summary of *N. lecontei* AMP orthology with other Hymenoptera

**Supplementary table S7.** Immunity gene orthologs between *N. lecontei* and *D. melanogaster.*

**Supplementary table S8 (separate xlsx file).** Insect olfactory receptor, gustatory receptor, odorant binding protein, cytochrome P450, and antimicrobial protein gene family sizes.

**Supplementary References**

**Supplementary file S1 (separate fasta file).** Amino acid and cDNA sequences for all manually annotated genes.

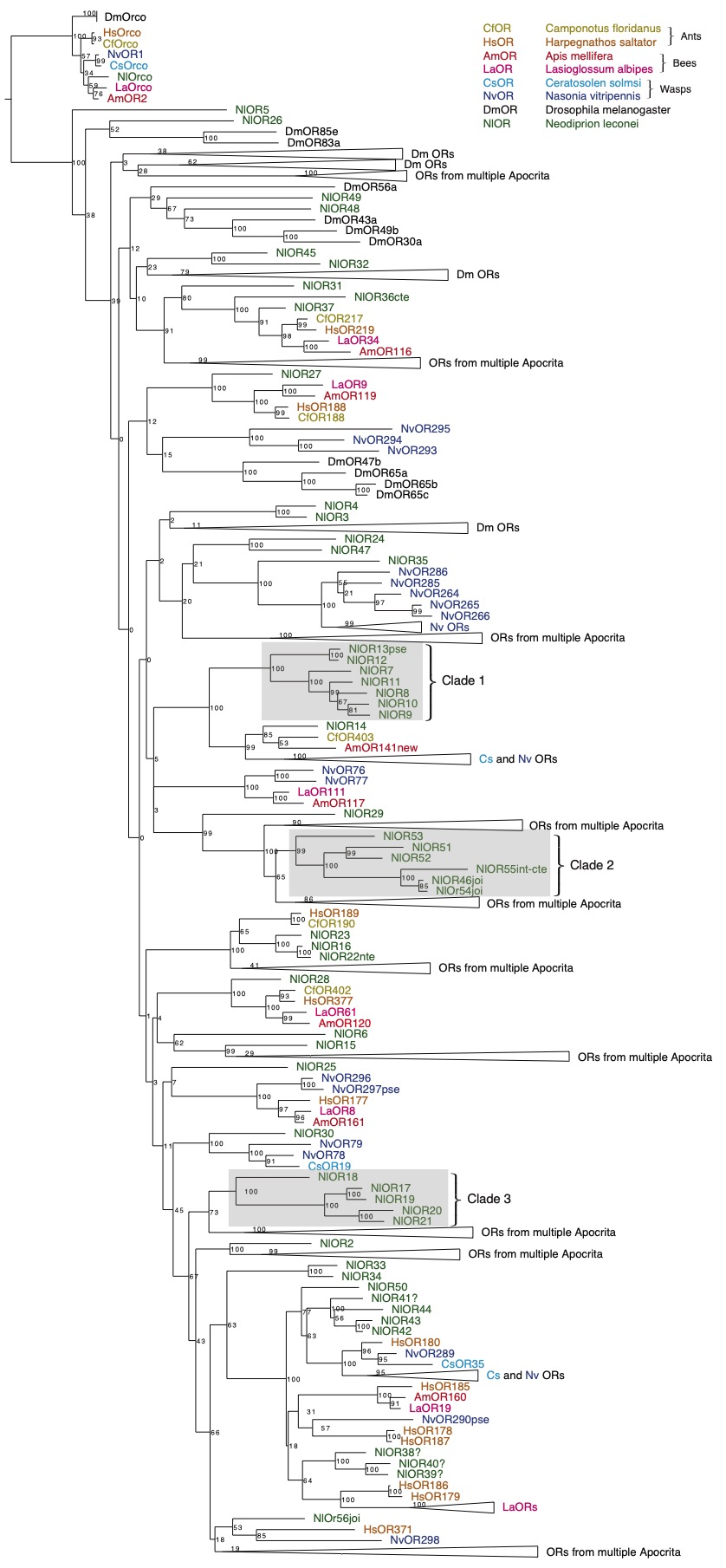

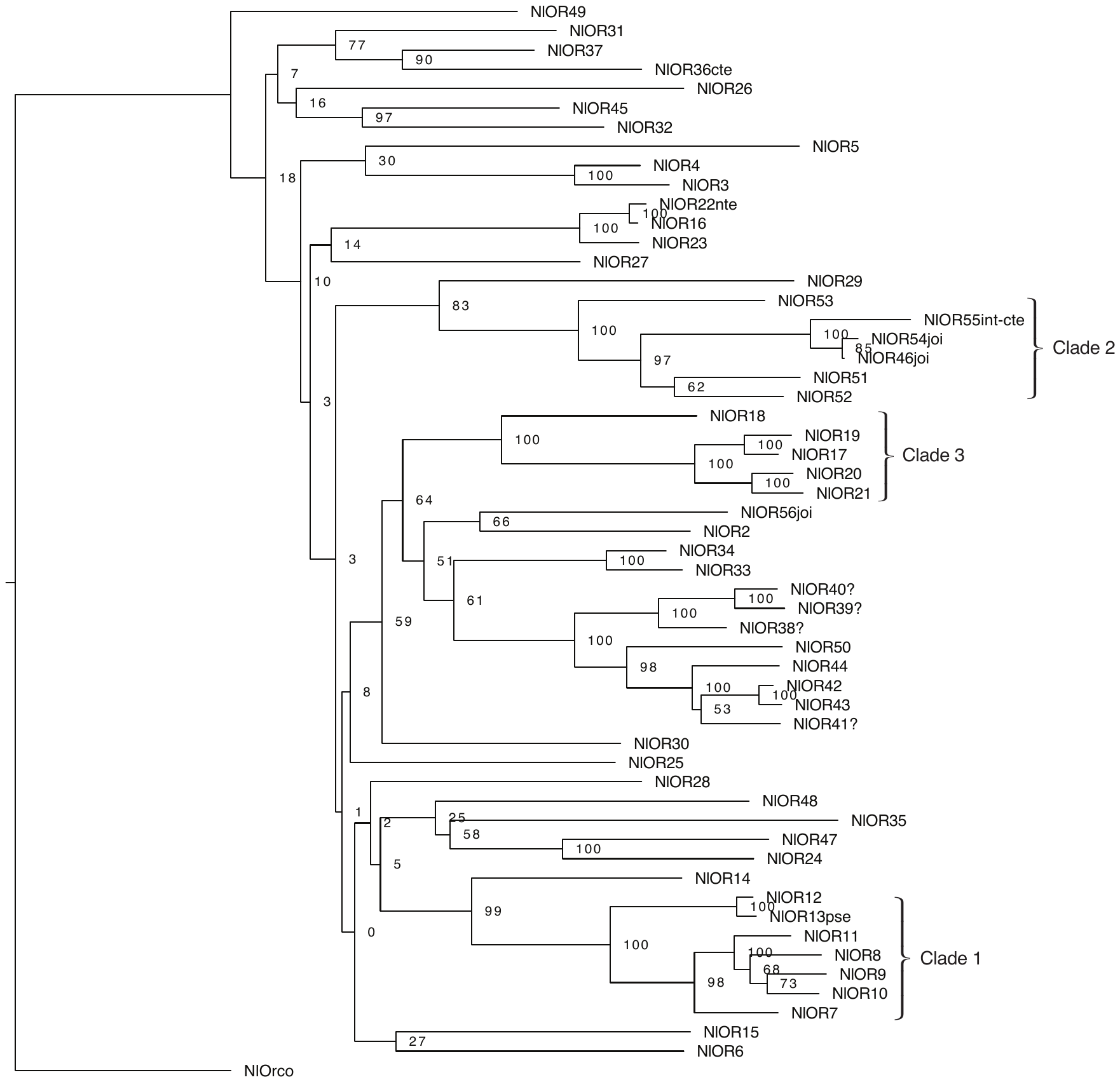

**Figure S1a. Olfactory receptor gene family amino acid phylogeny with select Hymenoptera.** Gray boxes indicate *N. lecontei* species-specific gene expansions that were tested for evidence of positive selection. Numbers are maximum likelihood bootstrap values. References: Robertson and Wanner (2006); Robertson et al. (2010); Smith et al. (2011); Zhou et al. (2012, 2015); FlyBase (accessed 25 Aug 17).

**Figure S1b. Olfactory receptor gene family cDNA phylogeny for *N. lecontei* only.** Clade names are the same as in Figure S1a.

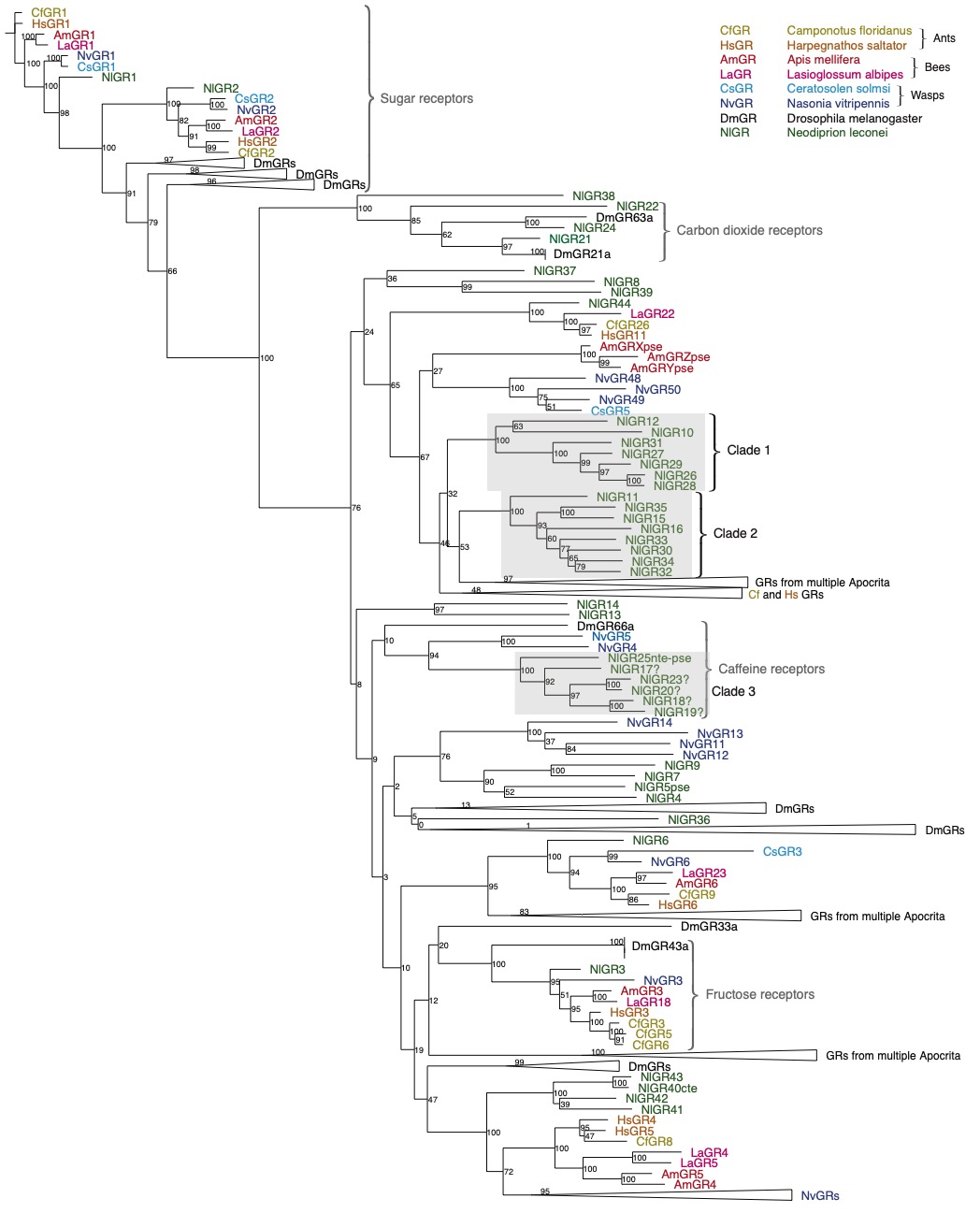

**Figure S2a. Gustatory receptor gene family amino acid phylogeny with select Hymenoptera.** Gray boxes indicate *N. lecontei* species-specific gene expansions that were tested for evidence of positive selection; numbers are ML bootstrap values. Known receptor functions (sugar, carbon dioxide, and caffeine receptors) are based on the *D. melanogaster* literature*.* References: Robertson and Wanner (2006); Robertson et al. (2010); Smith et al. (2011); Zhou et al. (2012, 2015); FlyBase (accessed 25 Aug 17).

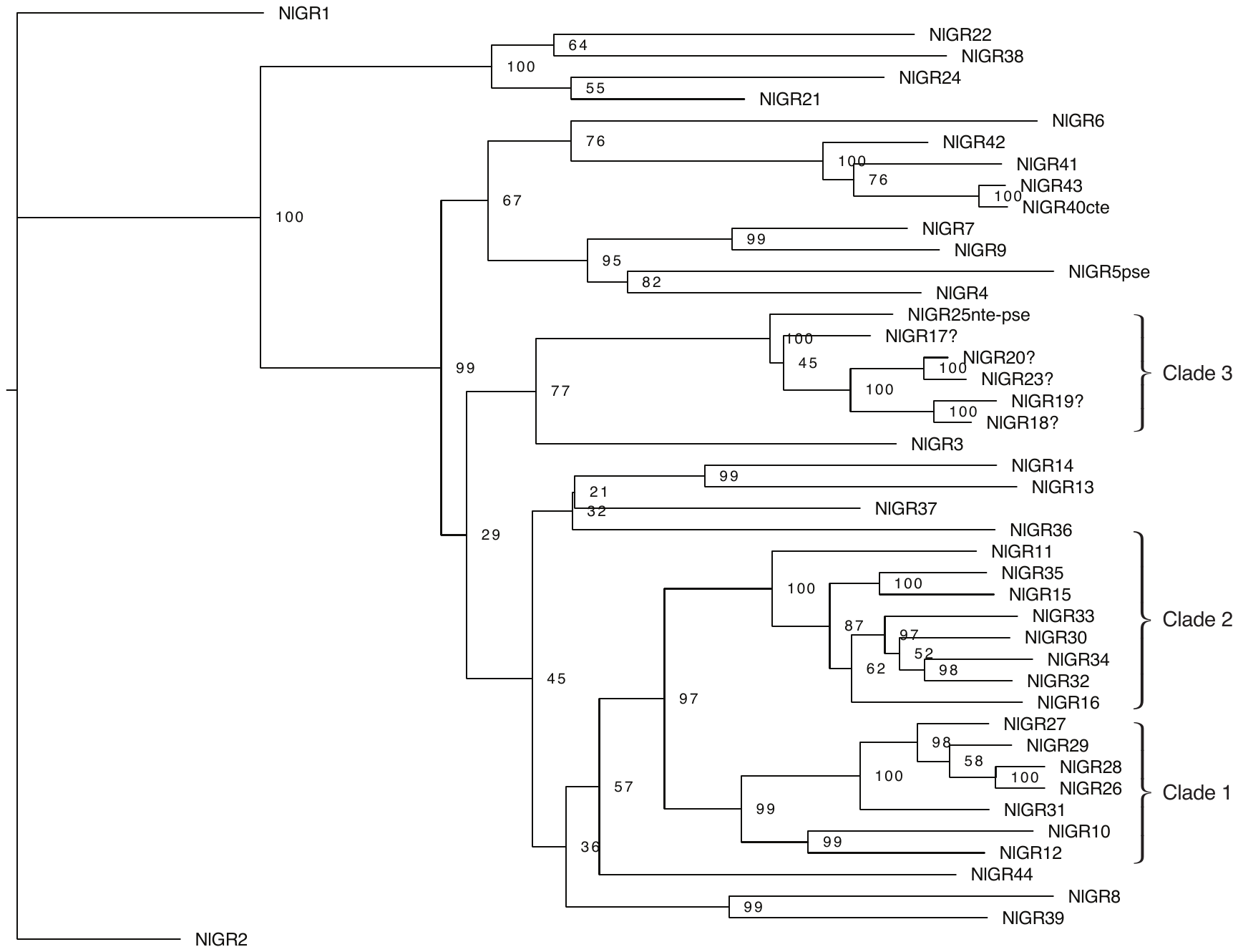

**Figure S2b. Gustatory receptor gene family cDNA phylogeny for *N. lecontei* only.** Clade names are the same as in Figure S2a.

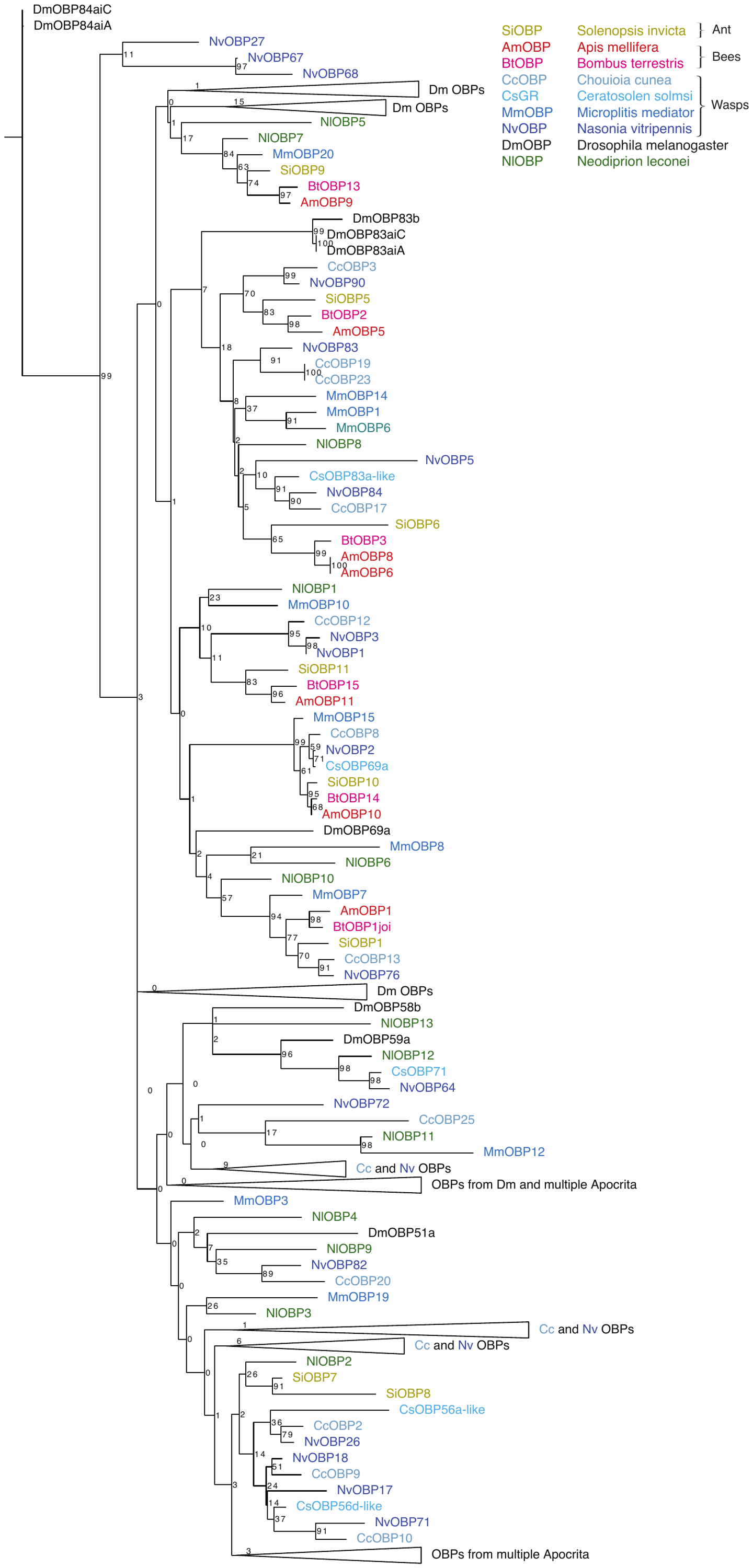

**Figure S3a. Odorant binding protein gene family amino acid phylogeny with select Hymenoptera.** Numbers are ML bootstrap value. No species-specific clades with ≥5 intact *N. lecontei* genes and a bootstrap score ≥70 were found. References: Forêt and Maleszka (2006); Gotzek et al. (2011); Vieira et al. (2012); Xiao et al. (2013); Sadd et al. (2015); Zhou et al. (2015); Zhao et al. 2016; FlyBase (accessed 25 Aug 17).

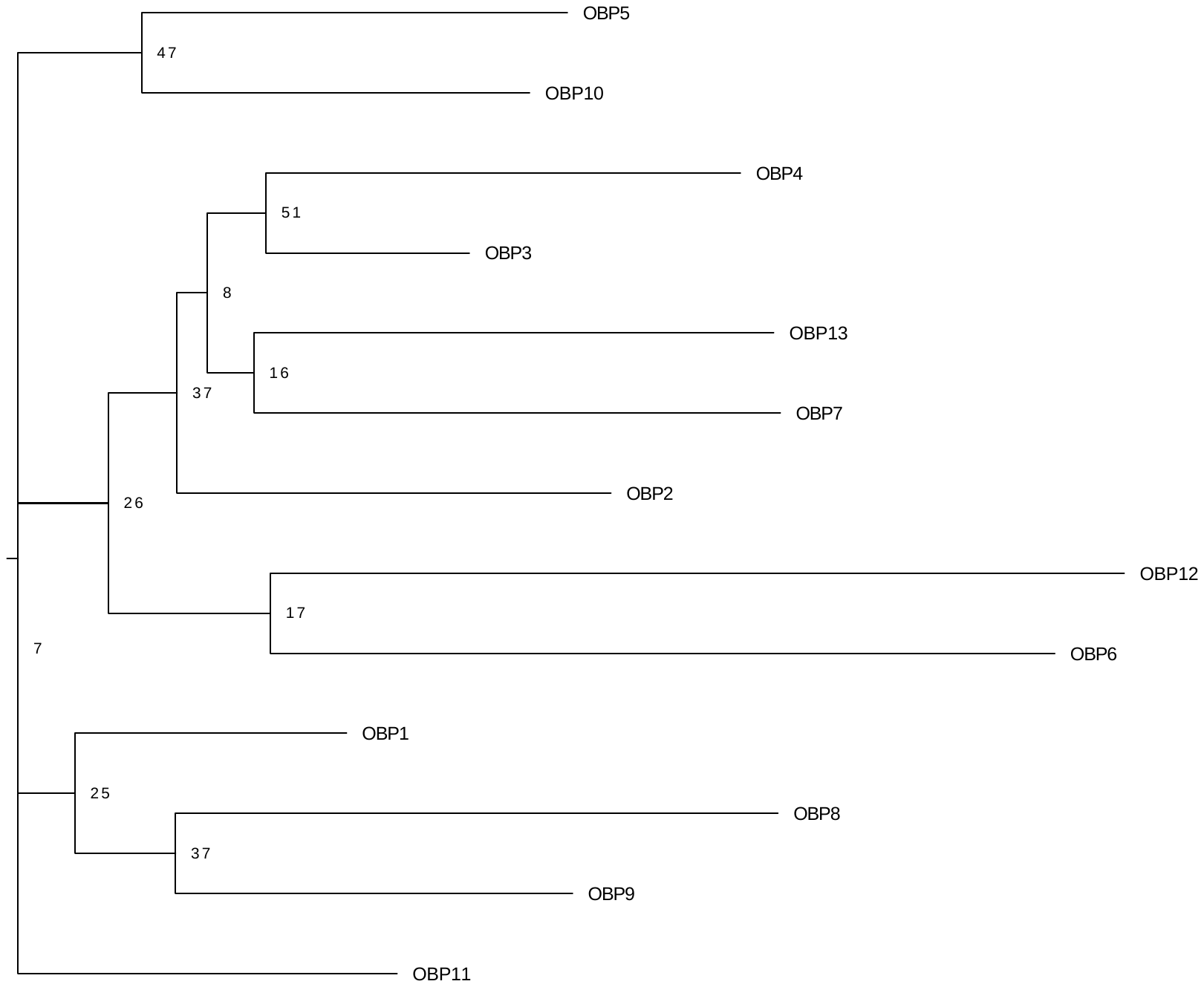

**Figure S3b. Odorant binding protein gene family cDNA phylogeny for *N. lecontei* only.**

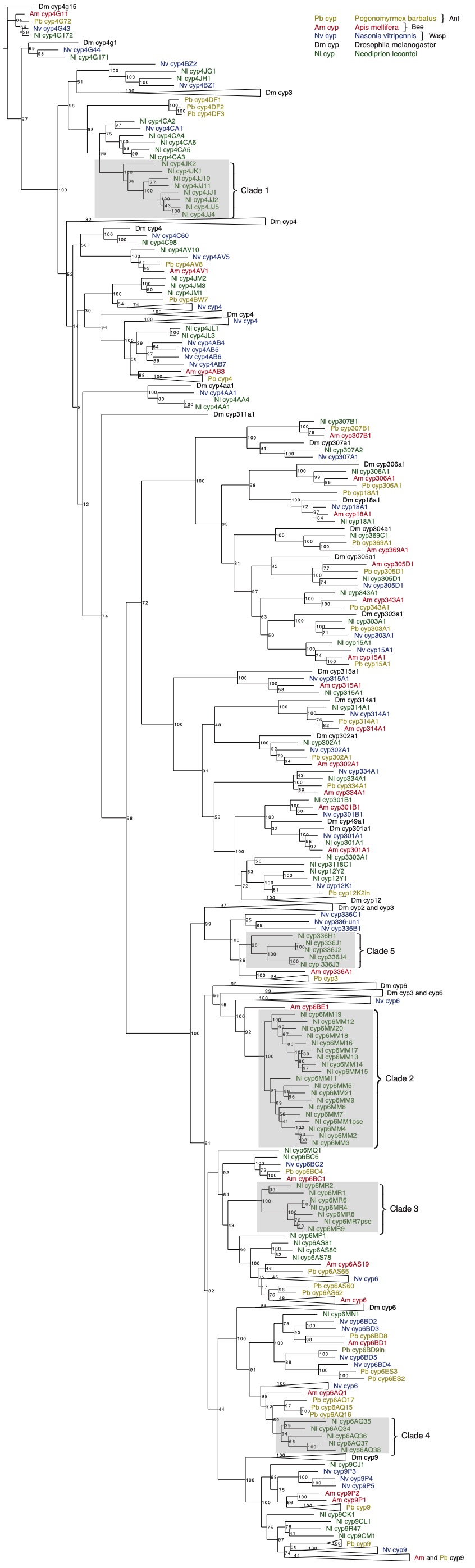

**Figure S4a. Cytochrome P450 gene family amino acid phylogeny with select Hymenoptera.** Gray boxes indicate *N. lecontei* species-specific gene expansions that were tested for evidence of positive selection. Numbers are maximum likelihood bootstrap values. References: Claudianos et al. (2006); Oakeshott et al. (2010); Smith et al. (2011); Feyereisen (2012).

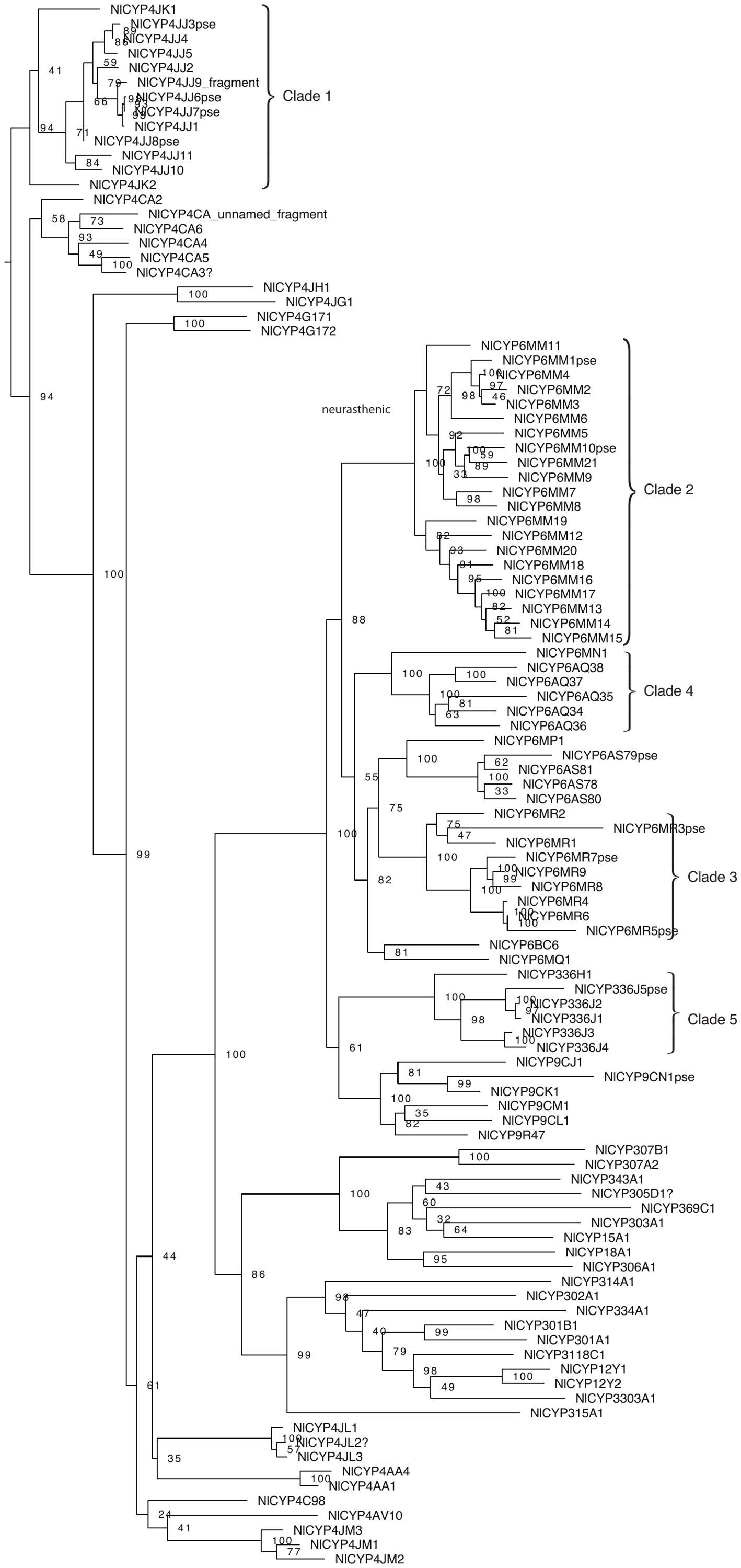

**Figure S4b. Cytochrome P450 gene family cDNA phylogeny for *N. lecontei* only.** Clade names are the same as in Figure S4a.

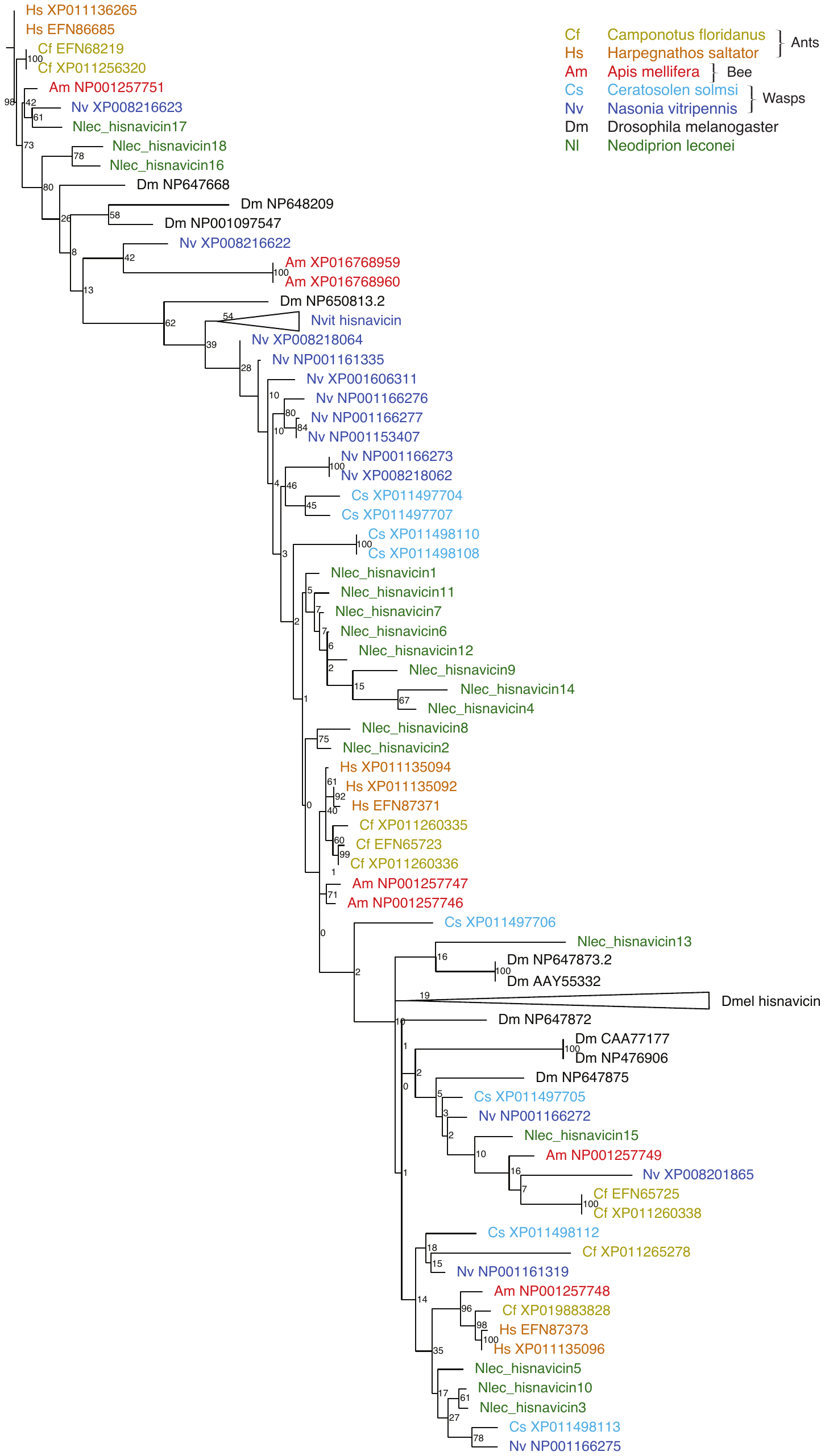

**Figure S5a. Hisnavicin gene family amino acid phylogeny with select Hymenoptera.** Although one species-specific clade had eight genes, it did not meet our requirements for a species-specific expansion because it had a bootstrap score <70. References: Evens et al. (2006); Lemaitre and Hoffmann (2007); Tial et al. (2010); Xiao et al. (2013); Gupta et al. (2015).

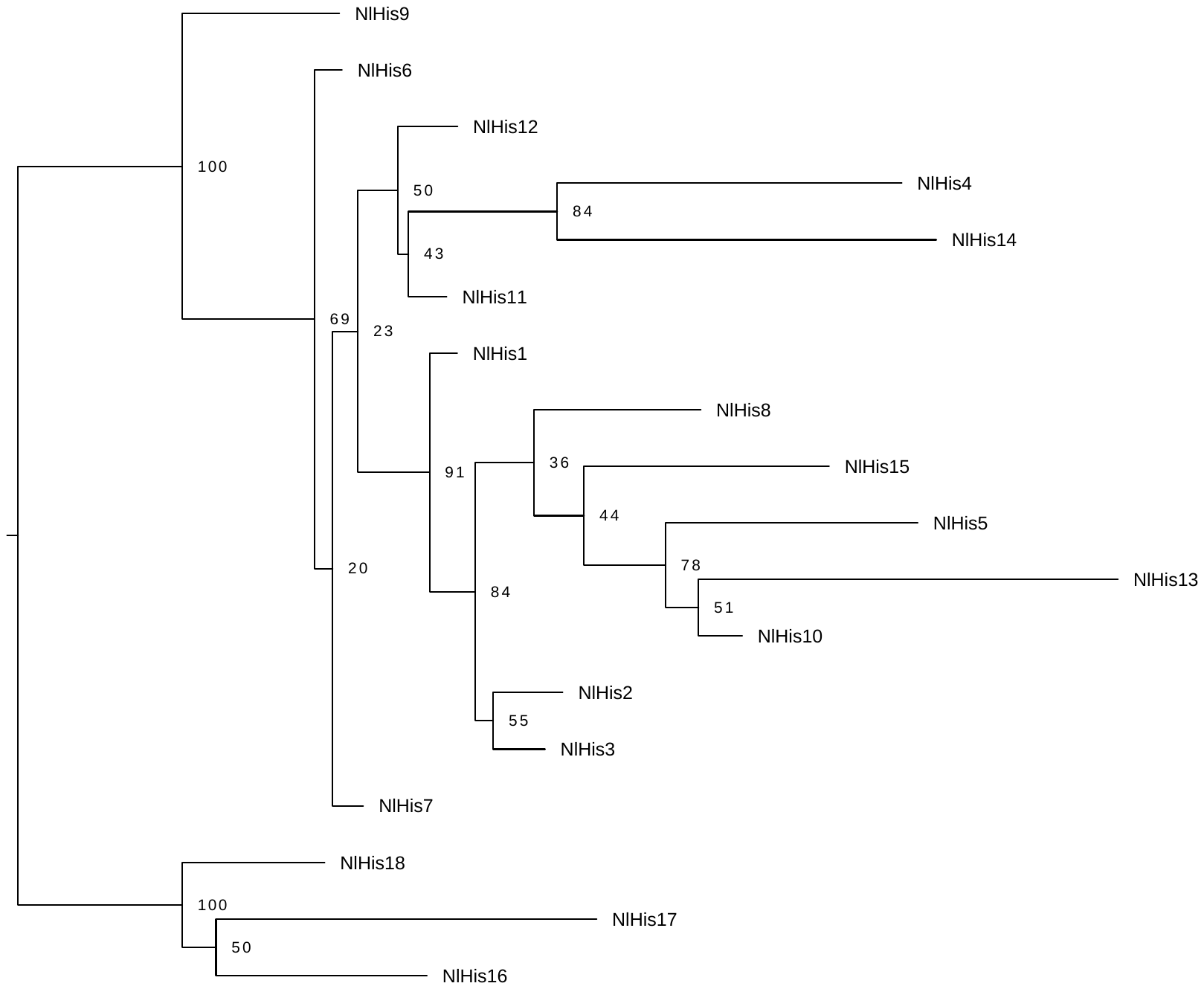

**Figure S5b. Hisnavicin gene family cDNA phylogeny for *N. lecontei* only.**

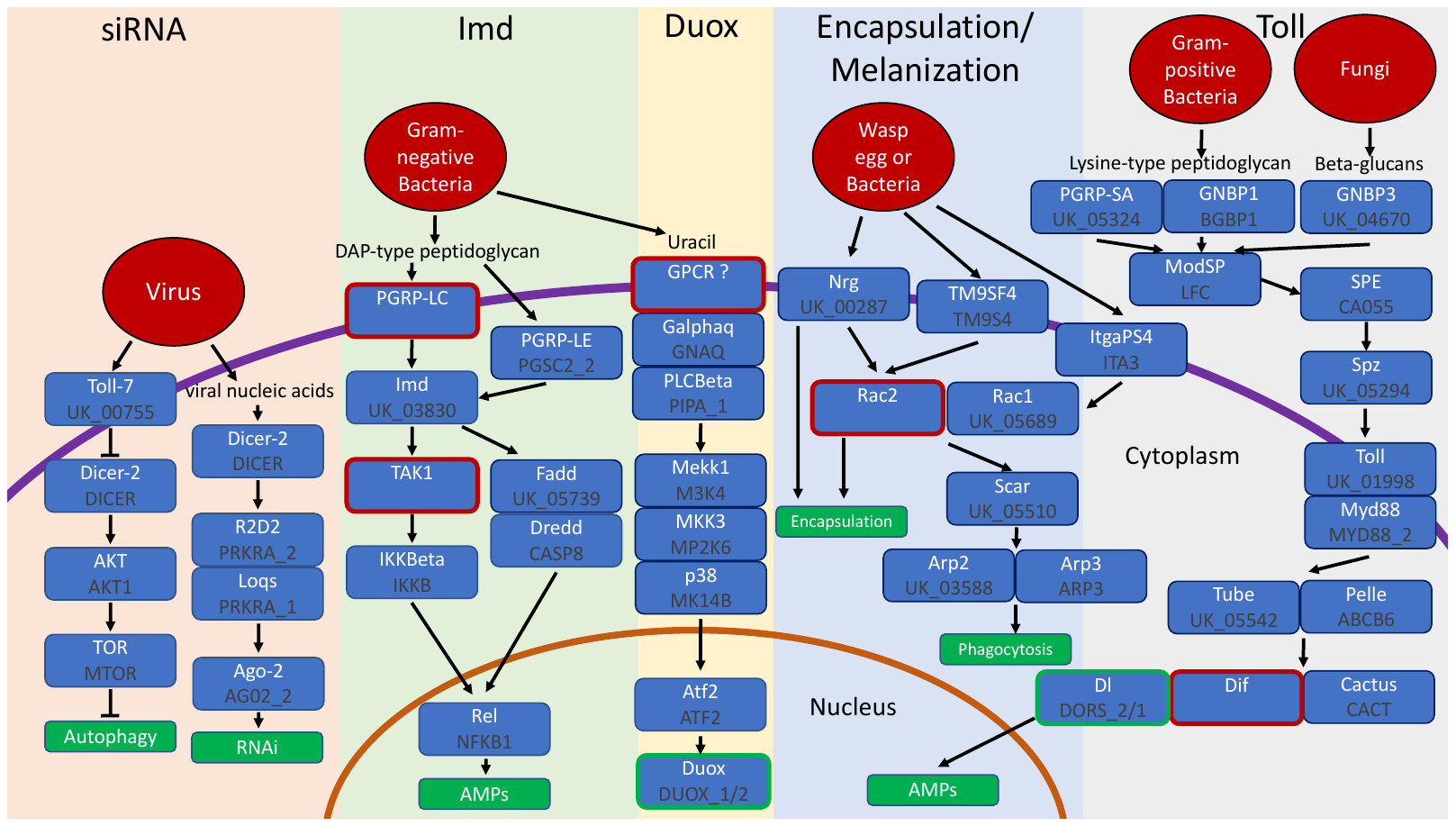

**Figure S6.** **An overview of *N. lecontei* immune pathways.** Gene names and pathways are based on *D. melanogaster* annotations. Red ovals represent immune challenge, blue boxes represent individual genes in each pathway, red outlines on blue boxes represent genes without one-to-one orthologs in *N. lecontei*, green outlines on blue boxes represent genes with more than one ortholog in *N. lecontei*, green boxes represent the end products of each pathway.

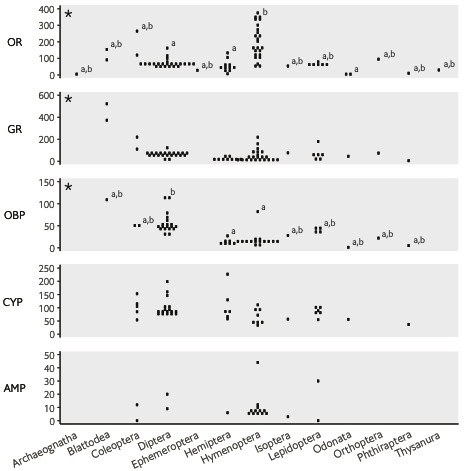

**Figure S7. Variation in gene family size among insect orders.** Each point represents the number of intact genes for a particular species belonging to that order. Asterisks indicate families for which there was significant size variation (*p* < 0.05 in Kruskal-Wallis rank-sum tests); letters indicate taxa that were significantly different after See Table S8 for references.

**Table S1.** Sequencing libraries and read counts for the *N. lecontei* genome

| Library type | Raw reads (PE) | Raw read depth | Filtered reads  (PE) | Filtered read depth |
| --- | --- | --- | --- | --- |
| 337 bp small-insert | 156,465,022 | 85x | 128,902,479 | 70x |
| 864 bp small-insert | 81,874,705 | 35x | 49,953,962 | 17x |
| 4.6 kbp mate-pair | 110,199,710 | 28x | 88,796,403 | 25x |

**Table S2.** Comparison of genome assembly and annotation statistics from published hymenopteran draft genomes.

| Species | Assembly version* | Genome size (Mbp)† | Assembly length (Mbp)‡ | Scaffold N50 (Mbp) | Total scaffolds | Total genes in OGS | GC content (%) | Repeat elements (%) | References |
| --- | --- | --- | --- | --- | --- | --- | --- | --- | --- |
| Sawflies |  |  |  |  |  |  |  |  |  |
| *Neodiprion lecontei* |  | 331 ± 10 (fc) | 239 | 0.243 | 4,523 | 12,980 | 40 | 16 | This paper |
| Bees |  |  |  |  |  |  |  |  |  |
| *Apis cerana* |  | 239 (?) | 228 | 1.4 | 2430 | 10,651 | 30 | 6 | Park et al. (2015) |
| *Apis mellifera* | Amel_4.5; OGSv3.2 | 262 ± 1 (fc) | 250 | 0.997 | 5,644 | 15,314 | 33 | 9 | Honeybee Genome Sequencing Consortium (2006); Elsik et al. (2014) |
| *Bombus impatiens* | Bimp_2.0 |  | 247 | 1.4 | 1,505 | 15,896 | 38 | 18 | Sadd et al. (2015) |
| *Bombus terrestris* | Bter_1.0 | 433 (fc) | 249 | 3.5 | 5,678 | 11,875 | 39 | 15 | Sadd et al. (2015); Stolle et al. (2011) |
| *Ceratina calcarata* |  | 364 (k) | 262 | 0.0736 | 3,577 | 11,310 | 42 |  | Rehan et al. (2016) |
| *Dufourea novaeangliae* |  | 342 (fc); 291 (k) | 291 | 2.4 | 84,187 | 12,453 | 40 | 37 | Kapheim et al. (2015) |
| *Eufriesea mexicana* |  | 1,940 ± 42 (fc);1,000 (k) | 557 | 0.0024 | 3,522,543 | 12,022 | 41 | 49 | Kapheim et al. (2015) |
| *Euglossa dilemma* |  | 3,200 (k) | 588 | 0.144 | 22,698 | 15,904 | 40 | 39 | Brand et al. (2017) |
| *Habropoda laboriosa* |  | 377 (k) | 294 | 1.3 | 650,185 | 13,279 | 39 | 26 | Kapheim et al. (2015) |
| *Lasioglossum albipes* |  | 416 (k) | 342 | 0.616 | 41,433 | 13,448 | 42 | 33 | Kocher et al. (2013) |
| *Megachile rotundata* |  | 273 (k) | 273 | 1.7 | 6,266 | 12,770 | 37 | 43 | Kapheim et al. (2015) |
| *Melipona quadrifasciata* |  | 257 (k) | 257 | 1.9 | 7,386 | 15,368 | 39 | 18 | Kapheim et al. (2015) |
| Ants |  |  |  |  |  |  |  |  |  |
| *Acromyrmex echinatior* |  | 335 (fc) | 300 | 1.1 | 16,221 | 17,278 | 34 | 28 | Nygaard et al. (2011) |
| *Atta cephalotes* | OGSv1.2 | 300 ± 1 (fc) | 318 | 5.2 | 2,835 | 18,093 | 33 | 25 | Tsutsui et al. (2008); Suen et al. (2011) |
| *Cardiocondyla obscurior* | Cobs_1.4 |  | 188 | 3.1 | 1,854 | 17,552 | 40 | 21 | Schrader et al. (2014) |
| *Camponotus floridanus* | Cflo_v4 | 238 (qPCR) | 238 | 0.603 | 25,494 | 17,064 | 34 | 15 | Bonasio et al. (2010) |
| *Cerapachys biroi* | Cbir_v3; OGSv1.8 |  | 214 | 1.3 | 4,579 | 17,263 | 42 | 14 | Oxley et al. (2014) |
| *Harpegnathos saltator* | Hsal_v3 | 297 (qPCR) | 297 | 0.598 | 21,347 | 18,564 | 45 | 27 | Bonasio et al. (2010) |
| *Linepithema humile* | OGSv1.1 | 251 ± 2 (fc) | 220 | 1.4 | 3,030 | 16,123 | 38 | 31 | Tsutsui et al. (2008); Smith CD et al. (2011) |
| *Pogonomyrmex barbatus* | OGSv1.1 |  | 235 | 0.793 | 4,646 | 17,177 | 37 | 9-18 | Smith CR et al. (2011) |
| *Pseudomyrmex gracilis* |  | 392 (k) | 282 | 0.35 | 6,556 | 16,069 | 39 | 40 | Rubin and Moreau (2016) |
| *Solenopsis invicta* |  | 463 (fd); 591(rk); 753.3 (fc) | 353 | 0.721 | 10,543 | 16,569 | 36 |  | Li and Heinz (2000); Johnston et al. (2004); Ardila‐Garcia et al. (2010); Wurm et al. (2011) |
| Wasps |  |  |  |  |  |  |  |  |  |
| *Nasonia vitripennis* | OGSv1.2 | 333 (fd) | 295 | 0.709 | 6,181 | 17,279 | 41 | 17 | Werren et al. (2010); Gregory (2013) |
| *Ceratosolen solmsi* |  | 294 (k) | 278 | 9.6 | 7,397 | 11,412 | 30 | 9 | Xiao et al. (2013) |

* Assembly and official gene set (OGS) version are v1.0 unless reported otherwise.

† Methods used to measure genome size: fc=flow cytometry, k=kmer analysis, fd=Feulgen densitometry, rk=reassociation kinetics, ?=unknown

‡ Scaffold length with gaps

**Table S3.** Summary of repetitive elements identified in the *N. lecontei* genome.

|  |  |  | **dnaPipeTE number of families** | **dnaPipeTE Percentage** |  | **RepeatModeler number of families** | **RepeatModeler Percentage (fraction of assembly)** | **RepeatModeler Percentage (fraction of total genome)** |  | **Portion of repetitive genome missing in assembly** |
| --- | --- | --- | --- | --- | --- | --- | --- | --- | --- | --- |
| **TE Order** |  |  |  |  |  |  |  |  |  |  |
| LTR |  |  | 64 | 0.032605 |  | 29 | 0.008601 | 0.006228 |  | 0.026377 |
| LINE |  |  | 62 | 0.009526 |  | 22 | 0.006415 | 0.004645 |  | 0.004881 |
| SINE |  |  | 29 | 0.000031 |  | 20 | 0.000042 | 0.000030 |  | 0.000000 |
| DNA |  |  | 63 | 0.038394 |  | 38 | 0.020358 | 0.014741 |  | 0.023653 |
| MITE |  |  | 0 | 0.000000 |  | 0 | 0.000000 | 0.000000 |  | 0.000000 |
| Helitron |  |  | 51 | 0.001384 |  | 32 | 0.001248 | 0.000903 |  | 0.000481 |
| **Repeat Order** | |  |  |  |  |  |  |  |  |  |
| rRNA |  |  | 21 | 0.007173 |  | 16 | 0.000173 | 0.000125 |  | 0.007048 |
| Low Complexity | |  | 8 | 0.006899 |  | 8 | 0.003830 | 0.002773 |  | 0.004126 |
| Satellite |  |  | 37 | 0.019220 |  | 21 | 0.003126 | 0.002263 |  | 0.016956 |
| Tandem repeats | |  | 0 | 0.000000 |  | 0 | 0.000000 | 0.000000 |  | 0.000000 |
| Simple repeat | |  | 8 | 0.037301 |  | 26 | 0.019386 | 0.014037 |  | 0.023264 |
| **Other** |  |  |  |  |  |  |  |  |  |  |
| Unknown |  |  | 24 | 0.123394 |  | 122 | 0.095414 | 0.069089 |  | 0.054305 |
| **Total** |  |  | 367 | 0.275928 |  | 334 | 0.158593 | 0.114837 |  | 0.161091 |

**Table S4.** Tissue types, read counts, and transcript counts for the *N. lecontei* transcriptome.

| Tissue | Raw MiSeq reads  (PE250) | Raw HiSeq reads  (PE50) | Filtered reads (combined) | Raw transcripts | Final transcripts |
| --- | --- | --- | --- | --- | --- |
| Adult female body | 6,915,054 | 34,695,138 | 32,163,481 | 26,527 | 26,335 |
| Adult female head | 2,577,645 | 34,479,620 | 31,528,864 | 28,577 | 28,386 |
| Adult male body | 2,580,520 | 37,763,308 | 34,496,435 | 25,754 | 25,644 |
| Adult male head | 3,181,430 | 29,909,654 | 27,442,849 | 30,884 | 30,775 |
| Feeding larval body | 1,624,852 | 32,351,773 | 29,142,629 | 22,139 | 22,092 |
| Feeding larval head | 3,798,521 | 34,720,778 | 31,521,407 | 28,253 | 28,114 |
| Eonymph body | 2,523,305 | 29,154,633 | 26,325,699 | 19,949 | 19,916 |

**Table S5.** Glomeruli counts from left and right antennal lobes of an adult *N. lecontei* male and an adult *N. lecontei* female.

| Sex | Side | Glomeruli |
| --- | --- | --- |
| Male | Left | 37 |
|  | Right | 37 |
| Female | Left | 49 |
|  | Right | 45 |

**Table S6.** Summary of *N. lecontei* AMP orthology with other Hymenoptera.

| **AMP Family** | ***Neodiprion lecontei* orthologs** | ***Nasonia vitripennis* ortholog** | ***Apis mellifera* ortholog** | ***Camponontus floridanus* ortholog** |
| --- | --- | --- | --- | --- |
| **Hymenoptaecins** | Nlec_unknown_03748-mRNA-1 | Nahymenoptaecin-1* (NP_001165829.1)  Nahymenoptaecin-2 (NP_001234886.1) | Hymenoptaecin (NP_001011613.1)** | Hymenoptaecin (XP_019883221.1) |
| **Abaecins** | Nlec_unknown_03749-mRNA-1 |  | Abaecin (NP_001011617.1) |  |
| **Tachystatin-type** | Nlec_unknown_04433-mRNA-3 | Naickin-1 (XP_016840398.1) |  | Cafickin1-1 (XP_011257508.1) |
| **Defensins** | Nlec_TEN2_1-mRNA-1 | Nasonin-2 (NP_001171933.1)*** |  |  |
| **Histidine-rich** | Nlec_CU21-mRNA-1  Nlec_PROML-mRNA-1 | Hisnavicin-4 (NP_001166274.1)  Hisnavicin-3  (NP_001166274.1)^ |  |  |

*reciprocal best Blast hit when more than one sequence is listed

**best blast hit for *A. mellifera* AMPs vs. *N. lecontei* proteins (not reciprocal)

***Note that while the best blast hit for nasonin-2 was Nlec_TEN2_1-mRNA-1, this was not a reciprocal best blast hit and Nlec_TEN2_1-mRNA-1 appears to be a teneurin.

^best blast hit for for *N. vitripennis* AMPs vs. *N. lecontei* proteins (not reciprocal)

**Table S7.** Immunity gene orthologs between *N. lecontei* and *D. melanogaster.*

|  | **Pathway** | **Gene** | **FBgn** | **Nlec ID** | **Orthology** | **%ID** |
| --- | --- | --- | --- | --- | --- | --- |
| **Viruses** | siRNA | *Toll-7* | FBgn0034476 | Nlec_unknown_00755-mRNA-1 | RBH | 53.1 |
|  |  | *PI3Kp2E* | FBgn0015279 | Nlec_PK3CB-mRNA-1 | RBH* | 45.5 |
|  |  | *Akt1* | FBgn0010379 | Nlec_AKT1-mRNA-1 | RBH | 72.4 |
|  |  | *Tor* | FBgn0021796 | Nlec_MTOR-mRNA-1 | RBH* | 61.1 |
|  |  | *Dicer-2* | FBgn0034246 | Nlec_DICER_mRNA-2 | RBH | 29.8 |
|  |  | *Loqs* | FBgn0032515 | Nlec_PRKRA_1-mRNA-1 | RBH* | 55.8 |
|  |  | *R2D2* | FBgn0031951 | Nlec_PRKRA_2-mRNA-1 | Dmel_v_Nlec | 28.0 |
|  |  | *Ago2* | FBgn0087035 | Nlec_AGO2_2-mRNA-1 | Dmel_v_Nlec** | 37.1 |
|  |  | *Drosha* | FBgn0026722 | Nlec_unknown_02096-mRNA-1 | RBH* | 70.2 |
| **Gram-negative** | Duox | *Duox* | FBgn0283531 | Nlec_DUOX_2-mRNA-1  Nlec_DUOX_1-mRNA-1 | RBH*** | 72.8  36.4 |
|  |  | *Galphaq* | FBgn0004435 | Nlec_GNAQ-mRNA-1 | RBH | 74.2 |
|  |  | *PLCBeta* | FBgn0262738 | Nlec_PIPA_1-mRNA-1 | RBH | 65.9 |
|  |  | *Mekk1* | FBgn0024329 | Nlec_M3K4-mRNA-2 | RBH | 46.7 |
|  |  | *MKK3* | FBgn0261524 | Nlec_MP2K6-mRNA-1 | RBH | 65.7 |
|  |  | *p38b* | FBgn0024846 | Nlec_MK14B-mRNA-1 | RBH | 79.1 |
|  |  | *Atf2* | FBgn0265193 | Nlec_ATF2-mRNA-1 | RBH | 46.7 |
|  | Imd | *PGRP-LC* | FBgn0035976 | NA | NA | NA |
|  |  | *PGRP-LF* | FBgn0035977 | Nlec_PGRP-mRNA-2 | RBH | 40.9 |
|  |  | *PGRP-LE* | FBgn0030695 | Nlec_PGSC2_2-mRNA-4 | RBH | 47.2 |
|  |  | *Imd* | FBgn0013983 | Nlec_unknown_03830-mRNA-1 | RBH | 72.8 |
|  |  | *Tak1* | FBgn0026323 | NA | NA | NA |
|  |  | *IKKBeta* | FBgn0024222 | Nlec_IKKB-mRNA-2 | RBH | 34.9 |
|  |  | *Rel* | FBgn0014018 | Nlec_NFKB1-mRNA-2 | RBH | 37.2 |
|  |  | *Kenny* | FBgn0041205 | NA | NA | NA |
|  |  | *Fadd* | FBgn0038928 | Nlec_unknown_05739-mRNA-1 | RBH**** | 28.0 |
|  |  | *Dredd* | FBgn0020381 | Nlec_CASP8-mRNA-1 | RBH | 26.1 |
| **Gram-positive** | Toll | *PGRP-SA* | FBgn0030310 | Nlec_unknown_05324-mRNA-1 | RBH | 42.2 |
|  |  | *GNBP1* | FBgn0040323 | Nlec_BGBP1-mRNA-4 | Nlec_v_Dnel | 32.2 |
|  |  | *GNBP3* | FBgn0040321 | Nlec_unknown_04670-mRNA-1 | RBH | 33.2 |
|  |  | *Psh* | FBgn0030926 | Nlec_unknown_06133-mRNA-1 | Dmel_v_Nlec | 32.0 |
|  |  | *modSP* | FBgn0051217 | Nlec_LFC-mRNA-1 | RBH | 28.1 |
|  |  | *SPE* | FBgn0039102 | Nlec_CA055-mRNA-4 | RBH***** | 38.5 |
|  |  | *Spz* | FBgn0003495 | Nlec_unknown_05294-mRNA-1 | RBH | 30.2 |
|  |  | *Toll* | FBgn0262473 | Nlec_unknown_01998-mRNA-1 | RBH | 35.2 |
|  |  | *Myd88* | FBgn0033402 | Nlec_MYD88_2-mRNA-2 | RBH*** | 37.9 |
|  |  | *Tub* | FBgn0003882 | Nlec_unknown_05542-mRNA-1 | RBH | 33.8 |
|  |  | *Pll* | FBgn0010441 | Nlec_ABCB6-mRNA-1 | Dmel_v_Nlec | 64.2 |
|  |  | *Dif* | FBgn0011274 | NA | NA | NA |
|  |  | *Dl* | FBgn0260632 | Nlec_DORS_2-mRNA-1  Nlec_DORS_1-mRNA-1 | RBH  Nlec_v_Dmel^a^ | 65.8  53.9 |
|  |  | *Cact* | FBgn0000250 | Nlec_CACT-mRNA-1 | RBH | 35.0 |
| **NA** | JAK/STAT | *Upd1* | FBgn0004956 | NA | NA | NA |
|  |  | *Upd2* | FBgn0030904 | NA | NA | NA |
|  |  | *Upd3* | FBgn0053542 | NA | NA | NA |
|  |  | *Dome* | FBgn0043903 | Nlec_unknown_01653-mRNA-1 | RBH^b^ | 26.5 |
|  |  | *Hop* | FBgn0004864 | Nlec_JAK-mRNA-1 | RBH | 42.5 |
|  |  | *Stat92E* | FBgn0016917 | Nlec_STA5B-mRNA-8 | RBH | 49.1 |
|  |  | *Ptp61D* | FBgn0267487 | Nlec_PTP61-mRNA-1 | RBH* | 59.5 |
|  |  | *RanBP3* | FBgn0039110 | Nlec_unknown_04013-mRNA-1 | RBH* | 35.0 |
|  |  | *Cnot4* | FBgn0051716 | Nlec_CNOT4-mRNA-1 | RBH* | 72.0 |
| **Cellular** | Encapsulation/ Phagocytosis | *Nrg* | FBgn0264975 | Nlec_unknown_00287-mRNA-1 | RBH | 60.4 |
|  |  | *TM9SF4* | FBgn0028541 | Nlec_TM9S4-mRNA-2 | RBH | 73.4 |
|  |  | *ItgaPS4* | FBgn0034005 | Nlec_ITA3-mRNA-1 | Dmel_v_Nlec* | 25.4 |
|  |  | *Mys* | FBgn0004657 | Nlec_unknown_04235-mRNA-1 | RBH | 55.2 |
|  |  | *Rac1* | FBgn0010333 | Nlec_unknown_05689-mRNA-1 | RBH | 92.7 |
|  |  | *Rac2* | FBgn0014011 | NA | NA | NA |
|  |  | *NimC1* | FBgn0259896 | NA | NA | NA |
|  |  | *Eater* | FBgn0243514 | Nlec_unknown_05111-mRNA-1 | RBH | 36.9 |
|  |  | *Scar* | FBgn0041781 | Nlec_unknown_05510-mRNA-1 | RBH* | 53.5 |
|  |  | *Arp2* | FBgn0011742 | Nlec_unknown_03588-mRNA-1 | RBH* | 87.7 |
|  |  | *Arp3* | FBgn0262716 | Nlec_ARP3-mRNA-1 | RBH* | 86.6 |
|  |  | *Tsr* | FBgn0011726 | Nlec_CADF_2-mRNA-1 | RBH* | 84.4 |
|  |  | *Chic* | FBgn0000308 | Nlec_PROF-mRNA-1 | RBH* | 80.2 |
|  |  | *Ssh* | FBgn0029157 | Nlec_SSH1-mRNA-1 | RBH* | 75.0 |
|  | Melanization | *PPO1* | FBgn0283437 | NA | NA | NA |
|  |  | *PPO2* | FBgn0033367 | Nlec_PRPA3_1-mRNA-1 | RBH^c^ | 60.1 |
|  |  | *PPO3* | FBgn0261363 | NA | NA | NA |

RBH = Reciprocal best hit

Dmel_v_Nlec = best hit for *D. melanogaster* protein against all *N. lecontei* proteins only

Nlec_v_Dmel = best hit for *N. lecontei* protein against all *D. melanogaster* proteins only

**Table S8.** Insect olfactory receptor, gustatory receptor, odorant binding protein, cytochrome P450, and antimicrobial protein gene family sizes. *Please see accompanying xlsx file. Tabs correspond to different gene families.*

**File S1.** Amino acid and cDNA sequences for all manually annotated genes. *Please see accompanying fasta file.*
